# Directing stem cell differentiation by chromatin state approximation

**DOI:** 10.1101/2025.04.24.650451

**Authors:** Luis F. Montano-Gutierrez, Sophie Müller, Ana P. Kutschat, Igor Adameyko, Davide Seruggia, Florian Halbritter

**Author notes:** Senior / corresponding authors or. Department of Neuroimmunology, Center for Brain Research, Medical University of Vienna, Vienna, Austria (Luis F. Montano-Gutierrez). Equal contributions (Luis F. Montano-Gutierrez, Sophie Müller).

## Abstract

A prime goal of regenerative medicine is to replace dysfunctional cells in the body. To design protocols for producing target cells in the laboratory, one may need to consider exponentially large combinations of culture components. Here, we investigated the potential of iteratively approximating the target phenotype by quantifying the distance between chromatin profiles (ATAC-seq) of differentiating cells *in vitro* and their *in-vivo* counterparts. We tested this approach on the well-studied generation of erythroblasts from haematopoietic stem cells, evaluating a fixed number of components over two sequential differentiation rounds (8×8 protocols). We found that the most erythroblast-like cells upon the first round yielded the most erythroblast-like cells at the second round, suggesting that greedy selection by chromatin approximation can be a viable optimisation strategy. Furthermore, by analysing regulatory sequences in incompletely reprogrammed chromatin regions, we uncovered transcriptional regulators linked to roadblocks in differentiation and made a data-driven selection of ligands that further improved erythropoiesis. In future, our methodology can help craft notoriously difficult cells in vitro, such as B cells.

## INTRODUCTION

The overarching goal of regenerative medicine is to replace diseased or damaged tissues with cells generated in a laboratory (1, 2). To minimise risks and maximize benefits, it is important to produce cells that are phenotypically equivalent to their *in-vivo* counterparts (1, 3). Moreover, cell production should be reproducible across laboratories and have a high yield (2, 4). A promising strategy is to direct the differentiation of stem cells using different types of external stimuli (chemical, physical) or genetic perturbations (5, 6). However, some cell types remain difficult to engineer in sufficient quality or quantity due to incomplete acquisition of the target phenotype, persistence of progenitor traits, or heterogeneity in the generated cell populations (3, 7). Moreover, cell culture components can be expensive and differentiation yields low, rendering cell production economically burdensome (8). Therefore, strategies to optimise, standardise, and reduce the cost of cell differentiation are highly sought after.

There are at least two technical challenges to the design of differentiation protocols. First, the number of culture components, concentrations, and application times is large and with each additional variation considered the number of possible protocols becomes exponentially larger. Without automation and multiplexed readouts, exhaustive assessment of all alternatives quickly becomes infeasible for most laboratories (9–13). Alternatively, the number of combinations to be evaluated may be reduced by experimental design, e.g., by applying mathematical modelling or engineering-inspired methodologies such as the Taguchi method for robust parameter design (13–18). These complex designs can be difficult to implement without bespoke hardware(19) and expertise, thus leading us to favour simple approaches. Borrowing a concept from machine learning (20), a “greedy” optimisation strategy may be an appealing solution to this problem. Greedy optimisation evaluates a fixed number of alternatives at each step and picks the option that is closest to the target as a basis for subsequent iterations. Implementing this approach to protocol optimisation could decrease the experimental burden by reducing the complexity of the search problem from exponential to a linear one, which has been useful to refine differentiation strategies in primate stem cells (21). However, if intermediate measurements are misleading, greedy selections may yield suboptimal solutions. The second technical challenge is how to assess the acquisition of the desired target cell identity. The typical approach is to evaluate assorted molecular markers via imaging, flow cytometry, or RT-qPCR. These assays are inexpensive and readily accessible at most institutions but provide only an incomplete assessment of the underlying phenotype, especially at intermediary differentiation stages. Transcriptomics assays have been widely adopted to assess phenotypes during differentiation (22–25). Combined with systems biology and network analysis, this allows not only to accurately assess the fidelity of differentiated cell types (3, 7, 26), but also to identify lineage-determinant transcription factors and signalling events (27–30) and thus a data-driven pre-selection of potential treatment targets for differentiation protocols (31). In practice, gene expression may be affected by circumstantial effects like proliferation state or environment, thus distracting from the key phenotype to be examined and contributing to so-called “batch effects”. In contrast, epigenomic features like DNA methylation, chromatin accessibility, and chromatin conformation are cell type-specific and can be measured effectively using genome-wide assays (e.g., ATAC-seq) (32–35). Previous work demonstrated that chromatin accessibility profiles are highly informative about cell types (33), that changes may even be detected earlier than at the RNA level while being stable against transient fluctuations, and that analysis of chromatin allows identifying cell-state-defining regulatory elements (36–40). The global epigenomic similarity may thus be a good proxy for the convergence toward a desired target cell type.

Here, we apply greedy optimisation logic to cell differentiation, using the haematopoietic system as a proof of principle. We take advantage of the iterative, multi-step nature of differentiation protocols, and use flow cytometry and ATAC-seq to quantify the phenotypic distance between the differentiating cells and the target, before the next iteration. By subjecting haematopoietic stem cells (HSCs) to two rounds of eight treatments each to produce erythroblasts (Ery), we show that the winning condition in the first round predicted the winners of the second round. Our results suggest that erythropoietin alone produces the best-matching erythroblasts, and that insulin (INS), heparin (HEP), and hydrocortisone (HYDRO), used in several protocols, may have an adverse, time-sensitive impact. Using machine learning, we reveal spurious accessibility of chromatin at binding sites of transcription factors such as STAT1, ZNF394, and IRF1/3, which are normally closed in reference erythroblasts. Finally, we find data-driven perturbations to target the activity of such transcription factors in future experiments. To facilitate the optimisation of other differentiation protocols, we provide a computational pipeline that performs the analysis and generates a report for test samples in comparison to a defined target cell type.

## RESULTS

We sought to evaluate the potential of greedy optimization for cell differentiation protocols by chromatin approximation using the example of erythroblast production from haematopoietic stem cells (HSCs). Published protocols for erythroid differentiation include hormones and cytokines such as erythropoietin (EPO), interleukin 3 (IL-3), insulin (INS), heparin (HEP), and hydrocortisone (HYDRO) (49, 50). Differentiation can be monitored based on loss of stem cell marker CD34, a gain and subsequent decline of CD36 (thrombospondin receptor) and CD71 (transferrin receptor), and finally a gain of CD235a (glycophorin A). As a baseline for treatment exploration, we took a well-established protocol (51) which consists of three rounds of treatments (hereafter referred to as Round 1-3; **Fig. 1a**). We focused on variations of three culture components in the first two rounds (followed my maturation in Round 3): the concentration of EPO (1x, 3x), the combined presence/absence of INS+HEP, and the presence/absence of HYDRO, yielding eight ingredient combinations per round, and 72 treatments in total (8 at end of round 1, plus 64 at round 2). Thus, by evaluating all treatments, we sought to learn if an optimum at round 1 (the greedy choice), on its own would lead to the global optimum (the best overall) at round 2. Throughout this manuscript we indicate the round of each treatment with the prefixes “1.” and “2.”. For example, addition of insulin and heparin to round 2 is indicated as 2.INS+HEP.

**Figure 1.**
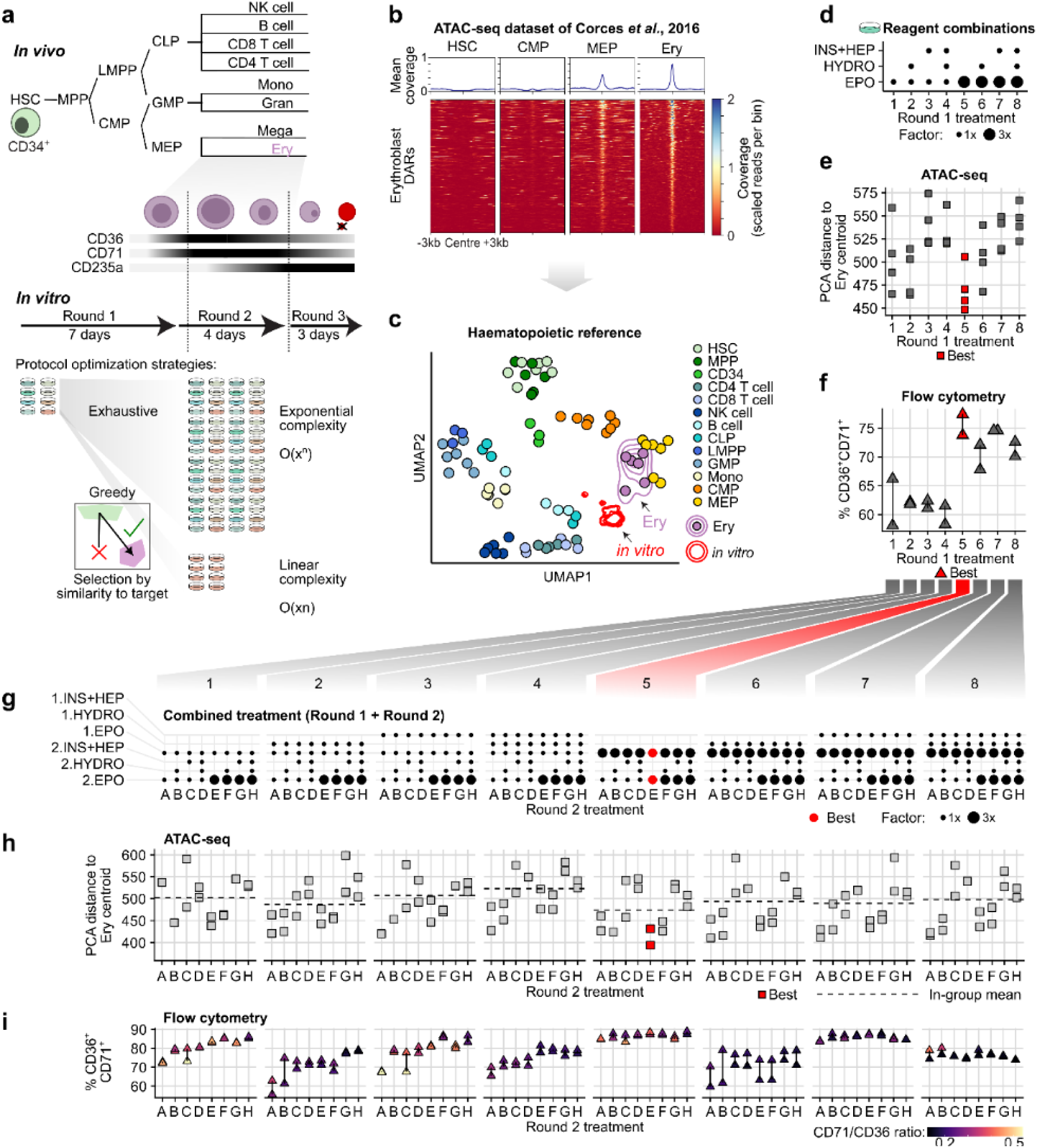
An exhaustive comparison of erythroid differentiation protocols confirms validity of greedy prioritization by chromatin similarity to approximate optimal solution. **a**, Top: Blood cells originate from CD34-positive haematopoietic stem cells (HSCs), which branch into lymphoid and myeloid lineages. Erythroblasts originate from Common Myeloid progenitors (CMP) and are marked by CD36, CD71, and CD235a expression. NK, natural killer; MPP, multipotent progenitor; LMPP, lymphoid–primed multipotent progenitors; CMP, common myeloid progenitor; mono, monocyte; gran, granulocyte; mega, megakaryocyte; GMP, granulocyte-monocyte progenitor; MEP, megakaryocyte-erythroid progenitor. Bottom: Two strategies to find protocols that approximate a target cell type. Exhaustive strategies evaluate all options but are costly and their complexity is exponential. We test whether a stepwise, greedy selection can result in optimal outcomes with lower experimental burden. **b**, Sorted region heatmap (DeepTools (74)) depicting the opening of erythroblast (Ery)-linked chromatin along differentiation in haematopoietic cells from a reference dataset(32). The regions shown were selected by differential accessibility analysis in erythroblast samples compared to all other samples in the reference (two-sided moderated t-tests, DESeq2 (75); P-value ≤ 0.05). **c**, UMAP of the cell type chromatin reference using cell type-informative PCA components shown in Supplementary Figure S1. Each point represents one independent samples. Density hulls indicate all samples from *in vitro* protocols (both rounds 1 and 2) generated in this study using different erythroid differentiation protocols (red), and the erythroblast target cells (purple). **d**, Ingredient combinations (8 total) tested for round 1 of differentiation, as a guide for data plots in panels e and f. Circle size indicates the multiplication factor of a base concentration of each ingredient added (see Methods). Absence of a circle indicates 0. **e**, Euclidean distance of all round-1 samples to the centroid of the Ery samples (“erythroid distance”) using the cell type-informative PCs (Supplementary Fig. S1). Each square represents one ATAC-seq datasets (2 technical replicates from 2 independent experiments each). Red colour indicates the treatment with samples closest to the Ery centroid, i.e. smallest distance. The distance values and metadata are available in Supplementary Table S1. **f**, Fraction of CD36 and CD71 double-positive cells from all treatments in panel **c** assessed by flow cytometry (n = 2 independent experiments). Red colour indicates the sample with the highest overall number of double-positive cells. The flow cytometry results are available in Supplementary Table S2. **g**, For an exhaustive evaluation of treatments, each sample from panel **c** (treatments 1-8) was subjected to a second round of the same eight treatment combinations (treatments A-H), resulting in 64 total treatment combinations (1A, 1B, etc.). **h**, Euclidean distance of each round-2 sample to the Ery centroid, analogous to panel d (n = 2 independent experiments). Red colour indicates the treatment with samples overall closest to the Ery centroid. Black dashed lines indicate the average distance for all samples coming from the same round-1 treatment. **i**, Flow cytometry analysis of round-2 samples, analogous to panel e (n = 2 independent experiments). Fill colour indicates the CD71/CD36 ratio, which differs even between samples with similar amount of CD71+CD36+ cells.

### Projection to a haematopoietic cell type reference reveals sequential approximation of the erythroblast phenotype and allows selection of high-performing protocols

We started by defining the target cell type to be approximated at the molecular level as a baseline to quantify the fidelity of cells differentiated *in vitro*. Previous research provided chromatin accessibility maps that distinguish major haematopoietic cell types, thus providing a suitable ground-truth reference for this purpose (32) (**Fig. 1b**). A principal component analysis (PCA) of the consensus chromatin peaks among all reference samples revealed that information about cell identity was not evenly distributed across all principal components (PCs) of the variance (**Supplementary Fig. S1a**). For example, PC1 separated cell types less clearly than PC3. To focus on features with a high cell type signal-to-noise ratio, we trained a multi-class Random Forest classifier on all PCs to classify all reference cell types and selected the most important PCs for the classification task (**Supplementary Fig. S1b**). Performing the same classification with this shortlist of cell-type-informative components reduced the classification error rate from ∼25.6% to ∼11.5%. Most misclassifications came from developmentally related cell types such as HSCs and multi-potent progenitors (MPPs), and different T cell subtypes. Using cell-type-informative PCs, we built a UMAP to visually summarise the cell type information in these cell-type-informative PCs (**Fig. 1c**).

Having defined the haematopoietic PCA space, we aimed to find treatments that produced cells with the closest resemblance to the reference erythroblasts. To find a high-performing differentiation strategy, we evaluated chromatin accessibility using ATAC-seq in duplicates (**Supplementary Fig. S2a-d**; overview, metadata, and quality metrics of all genomics data: **Supplementary Table S1**) and the number of CD36+CD71+ double-positive cells using flow cytometry (**Supplementary Fig. S3a-b, Supplementary Table S2**). After confirming data were of satisfactory quality (average FRiP score = 0.24, **Supplementary Table S1, Supplementary Fig. S2a**) and reproduced expected chromatin accessibility patterns at the well-described *HBB*, but not the foetal and embryonal beta globin genes *HBG1/2* and *HBE* (**Supplementary Fig. S2b, c**), we visually inspected the results of the differentiation experiments by projecting the ATAC-seq data to the haematopoietic reference and observed that most treatments appear globally close to the reference erythroblasts (**Fig. 1c, Supplementary Fig. S1a**), with erythroblast-linked chromatin regions opening (**Supplementary Fig. S2d**). To evaluate treatment performance more objectively, we calculated the Euclidean distance in PCA space between each sample and the centroid of all erythroid samples. We reasoned that this distance provided a unidimensional, interpretable objective function to minimise as the samples converged towards the erythroid distribution. PCA distance comparisons revealed treatments that helped cells approach the average erythroid chromatin profile (i.e., they reduced the PCA distance), whereas others failed to match some Ery chromatin features and increased the distance (**Fig. 1e**). At the end of round 1, we identified samples from treatment 5 as the closest to the erythroblast centroid. This treatment, which consisted of only high EPO, also yielded the highest number of CD36+CD71+ double-positive cells (**Fig. 1f, Supplementary Fig. S3b**). This consistency suggests high concordance between conventional metrics of cell fidelity and our chromatin distance metric. Thus, according to the greedy search principles, treatment 5 would be the one elected forward for round 2.

To assess whether the greedy choice of treatment 5 would eventually yield the best erythroblast-like samples, we then evaluated all samples corresponding to differentiation round 2 (**Fig. 1g-i**). Treatments building on top of treatments 5 (5A-H) and 7 (7A-H) from round 1 produced the highest percentage of CD71+CD36+ cells (∼90%), however, the former had a higher CD71/CD36 ratio (**Fig. 1i, Supplementary Fig. S3c-d**), which suggests further maturation (52). Concordantly, treatments 5A-H also had a smaller PCA distance to the centroid of all erythroid samples (called “erythroid distance” henceforth) than 7A-H (mean distance 473.9 vs 489.1; **Fig. 1h**, black dashed lines). Treatment 5E (**Fig. 1h-i, Supplementary Fig. S3e, f;** highlighted in red), produced samples with a smaller erythroid distance than all others, thus representing the overall best treatment for erythroblast production in our hands.

We sought to test the robustness of our greedy selection to technical variation. To this end, we performed a computational simulation in which we varied the quality of the test and reference samples by down-sampling read counts assigned to each peak thus emulating effects of different sequencing depths. We found that erythroid distances calculated on down-sampled datasets were highly correlated with the original metric down to a sequencing depth of only ∼5M reads in the test dataset, if the reference was of a sufficient quality (≥10M reads; **Supplementary Fig. S4a-b**). With low depth in the reference dataset, results became unstable with a high variation between random trials (**Supplementary Fig. S4b**). This is due to a decreasing margin between the erythroid distance of the winning sample (= lowest erythroid distance) and all other samples (**Supplementary Fig. S4c**), resulting in a variable assignment of winning treatment across trials (**Supplementary Fig. S4d**). However, at ≥15M average depth of the test data, treatment 5 was consistently picked as the winner in all trials (**Supplementary Fig. S4e**). Treatment 5E was most frequently chosen as the winner of round 2, but at low to medium sequencing depths (<20M), other protocols were sometimes favoured including 7A (the second-best treatment in our main analysis).

Furthermore, we tested whether chromatin distances were informative in other differentiation lineages by reanalysing public ATAC-seq data from three *in-vitro* differentiation studies. First, we confirmed that the lowest erythroid distance correlated with grades of ongoing maturity defined by FACS markers in independently performed experiments based on independent protocols and other biological source materials (45) (**Supplementary Fig. S5a**). Second, we found that chromatin distance to monocytes in our reference atlas (“monocyte distance”) was lowest in HL60 cells differentiated using a monocytic differentiation protocol compared to those subjected to macrophage or neutrophil maturation (44), supporting that our concept for protocol selection might be generalisable to other lineages (**Supplementary Fig. S5b**). Third, we explored distance to NK cells in our references based on *in-vitro*-differentiated HSPCs (43). In this analysis, chromatin-based “NK cell distance” varied strongly with only a marginal decrease throughout differentiation (**Supplementary Fig. S5c**). NK, CD4+, and CD8+ T cells are similar at chromatin level (**Supplementary Fig. S1**), which might be a reason why chromatin distance is less informative in this instance.

Our results confirm that a greedy choice during the first round of differentiation could effectively lead to the overall best-performing treatment, and thus that greedy treatment searches are a viable way to optimise differentiation. However, caution is advised when applying the approach in other lineages, where the distinguishability of cell types in the reference might be one indicator of cases where chromatin distance is not a sufficient indicator of differentiation progress.

### Combined insulin and heparin induce time-dependent effects on chromatin during erythroid differentiation

To explain why some samples were closer to the erythroid centroid than others, we further scrutinised how the PCA distance was affected by the treatments by quantifying the effects of each ingredient on the distance metric via a mixed-effects model. Briefly, this approach explains variance in the chromatin distance as a sum of biases, each of them brought by one ingredient. These effects are allowed add onto each other or cancel each other out. Thus, the model estimates which linear coefficients for every ingredient would create a best fit for the observed distances and therefore explains treatment diversity in our data. Having all possible combinations of treatment components allows to interrogate the effects of each ingredient.

Upon modelling the effects of individual treatment components over 64 protocols, we found that EPO had the strongest positive effect on differentiation outcome (**Fig. 2a, Supplementary Fig. S6a**). According to the model, each unit of EPO reduced the distance to the erythroid centroid by a factor of ∼7.8 on average (**Supplementary Table S3**). In contrast, we found that incorporation of INS+HEP had a stark hindering effect, particularly in round 2 (2.INS.HEP); its addition increased the PCA distance by ∼30 points on average (**Fig. 2b**). Inspection of the data confirmed a significant association of 2.INS+HEP with large erythroid distance (p = 1.5 * 10^-15^, two-sided Wilcoxon rank sum test, P-values adjusted for multiple hypothesis testing using the Benjamini-Hochberg method), while 1.INS+HEP had only a marginal effect (**Fig. 2c, d**). That is, the effect of INS+HEP during the first differentiation round was eventually overwritten by the treatment in round two. In contrast, the subtle effect of high EPO in round 1 prevailed independent of ensuing treatment choices (**Supplementary Fig. S6a, b**). HYDRO had diverging effects at early and late stages, but neither of those effects were significant (**Fig. 2b, Supplementary Fig. S6c**).

**Figure 2.**
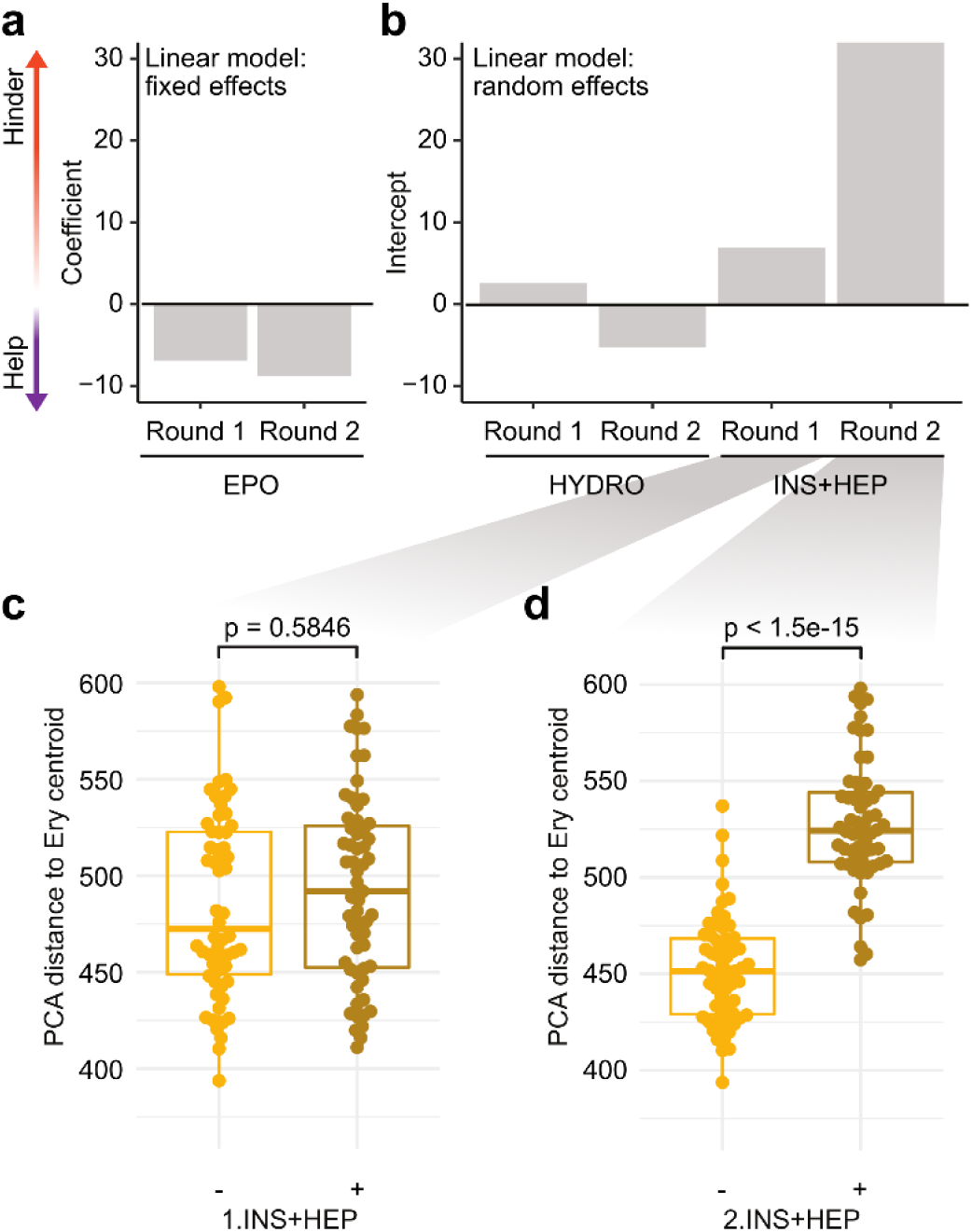
Statistical mixed-effects modelling identifies the impact of treatment ingredients on erythroid differentiation. **a,** We used a mixed effects model to quantify the relative impact of treatments on the PCA distance to the Ery centroid (cp. Fig. 1e, h). Fixed effects are Erythropoietin (EPO) units at both round 1 and round 2 of *in-vitro* differentiation. Negative and positive coefficients (left) indicate the estimated effect on erythroid distance per unit of treatment, respectively. The parameters of the model are available in Supplementary Table S3. **b,** Intercepts for variables included as random effects in the model described in panel a: insulin plus heparin (INS+HEP) and hydrocortisone (HYDRO) at round 1 and round 2. **c,** Box plot showing the median and interquartile range (IQR) of the distance to the erythroblast centroid for samples of differentiation round 2 (n = 119 samples), divided by whether they received INS+HEP during round 1. Whiskers extend to 1.5*IQR. All P-values correspond to a two-sided Wilcoxon rank sum test adjusted for multiple hypothesis testing using the Benjamini-Hochberg method. **d**, Boxplots as in panel c, showing samples divided by INS+HEP at round 2.

Altogether, this suggests that the timing of some of ingredients may impact differentiation performance and, more generally, that the distance to the target in the cell type-informative PCA can capture large systematic effects of treatments.

### In-vitro differentiated erythroblast-like cells show residual accessibility of progenitor chromatin modules

Our preceding analysis provided a bird’s-eye view of effects of treatments on the phenotype of differentiating cells. We wondered which genomic regions where most affected by the treatments at the chromatin level and how this influenced the acquisition of erythroblast identity. To investigate this question, we searched for chromatin regions whose accessibility would be highly predictive of a cell type, irrespective of how large its fold change was. To this end, we employed Receiver Operating Characteristic (ROC) analysis to evaluate 314, 431 high-priority regions (see **Methods**) on their ability to discriminate each cell type based on accessibility (**Supplementary Fig. S7a**). For instance, an ROC area under the curve (AUC) of 1 would indicate that all samples of a given cell type (and no other samples) had the highest accessibility and thus the region was unequivocally predictive for that cell type. An AUC ≤ 0.05 would indicate that the region was uniquely inaccessible in that cell type and thus was also (negatively) predictive. Lastly, an AUC of ∼0.5 is equivalent to a random guess between two groups.

In total, we found 35, 533 peaks with a ROC AUC greater or equal than 0.95 for any cell type, (**Fig. 3a, Supplementary Fig. S7b-c, Supplementary Table S4**). The selected regions were often proximal of genes expressed in the respective cell types (**Supplementary Fig. S7d, Supplementary Table S5**) and of DNA binding motifs of transcription factors (TFs) with a known association to the queried cell types (**Fig. 3c**, **Supplementary Table S6**). For example, motifs of the KLF-family TFs were exclusively enriched in erythroblast-specific regions, consistent with the key role of KLF1 in erythroid specification (53–55). This confirms that the ROC analysis identified biologically relevant regions, which we refer to as cell type-specific chromatin modules.

**Figure 3.**
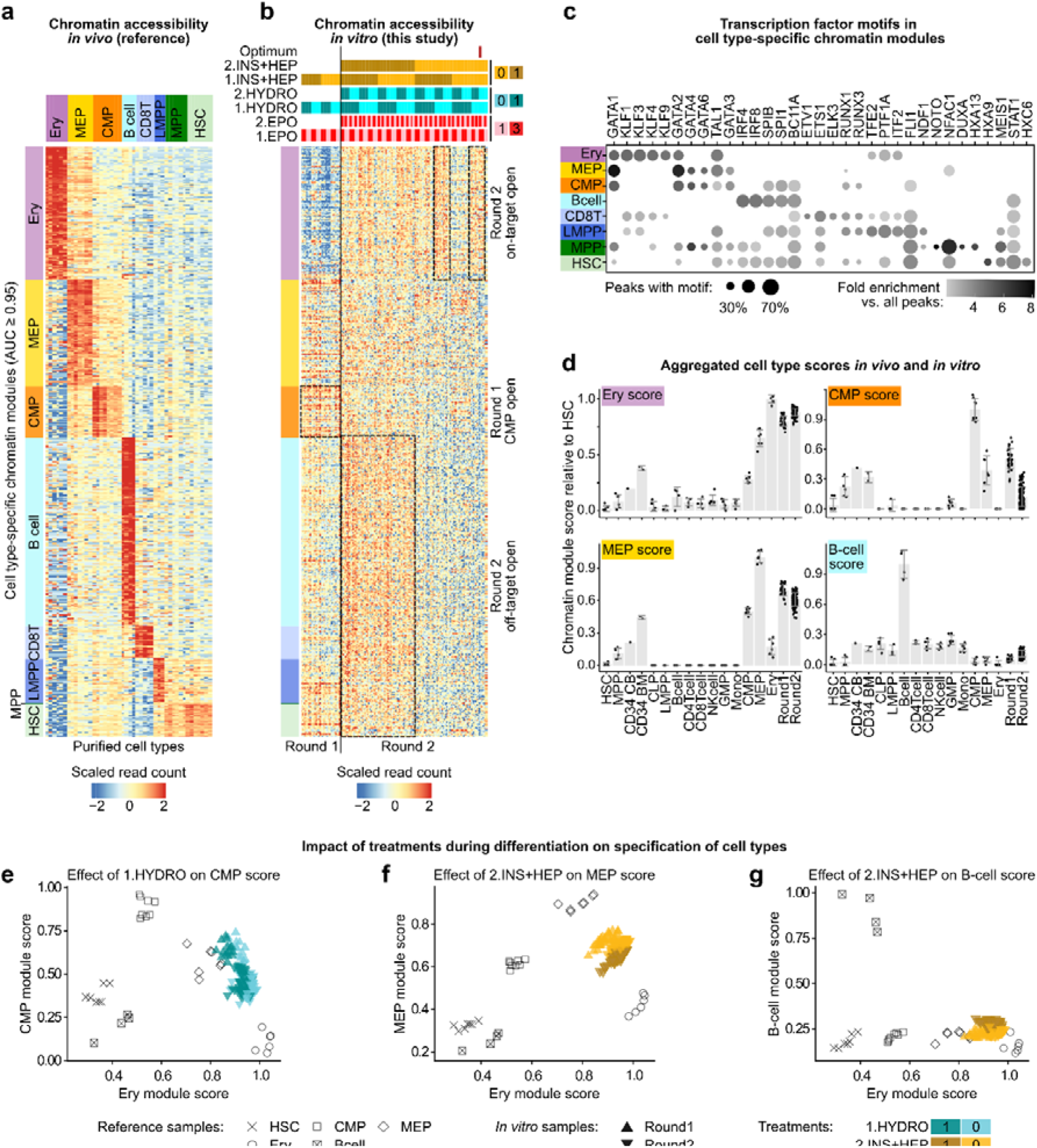
Effects of treatment ingredients on cell type-specific chromatin regions. **a**, Heatmap showing depth-normalised counts (row-scaled) of cell type-specific chromatin regions. Cell type-specific chromatin modules were defined by the area under the curve (AUC) of the receiver operating characteristic (ROC) to be strongly predictive of each cell type in the haematopoietic reference (32), see Supplementary Figure S7a. Ery, erythroblast; MEP, megakaryocyte-erythroid progenitor; CMP, common myeloid progenitor; LMPP, lymphoid-primed multipotent progenitor cells; MPP, multipotent progenitor; HSC, haematopoietic stem cell. Annotated peaks are provided in Supplementary Table S4. n(peaks) = 595, 523, n(samples) = 78. **b**, Heatmap analogous to panel a, showing the accessibility of samples from *in-vitro* differentiation generated in this study. The ingredient concentrations used for each sample are indicated on top. Selected blocks of coordinates chromatin changes have been highlighted manually with black boxes, as described in the text. n(samples) = 151. **c**, Enriched transcription factor (TF) binding motifs of each cell type-specific chromatin module relative to all consensus peaks found in the reference dataset. Regions with >0 motif hit for a TF were counted for all regions in each module (circle size), compared to the background (fold change indicated by colour). The top-5 enriched TFs were selected for visualization and all enrichment results (hypergeometric test) are listed in Supplementary Table S6. **d**, Barplots of relative module scores of the cell identity modules in panel a. Scores were calculated across all samples using single-sample gene set variation analysis (47) (GSVA). All scores in the bar plots were scaled such that a score of 0 matched the average of hematopoietic stem cells and 1 matched the average score for the original cell type (see Methods). Error bars correspond to the mean +/− the standard deviation for each subgroup. **e**, Biplots showing contrasting the effect of hydrocortisone (HYDRO) in round 1 on Ery module score and the CMP module score. Plotting symbols depict a different subgroup of samples, with symbols indicating the cell type and colour indicating the treatment status (for *in-vitro* samples only). n(samples) = 183. **f**, Biplot analogous to panel e showing the effect of insulin+heparin (INS+HEP) in round 2 on the Ery and MEP module scores. **g**, Biplot showing the effect of insulin+heparin (INS+HEP) in round 2 on the Ery and B-cell module scores.

Having confirmed that our collection of cell identity peaks constituted functional chromatin modules of each cell type, we then interrogated the activity of each module in our *in-vitro* samples (**Fig. 3b**). We found that treatments systematically influenced the accessibility of the modules. In line with our observation that INS+HEP in round 2 had a hindering effect on erythroid differentiation (cp. **Fig. 2**), samples with this treatment showed increased accessibility of non-Ery modules, including the HSC module, which suggests incomplete differentiation (**Fig. 3b**). Additionally, samples resulting from treatments using HYDRO at round 1 displayed low accessibility in the erythroblast-specific module **(Fig. 3b).**

These observations suggest that different treatments impacted chromatin beyond the cell type modules of erythroid cells. To grasp the relative magnitude of these effects, we used the accessibility of all regions in each cell type module to quantify chromatin-based “cell type module scores”, using single-sample gene set variation analysis(56) (**Fig. 3d)**. All reference cell type samples scored 1.0 on average in their own cell type score, while cell types of the same lineage (e.g., common myeloid [CMP], megakaryocyte-erythroid progenitors [MEP], erythroblasts) also had non-zero scores for related modules (**Fig. 3d**). The score approached ∼0 with more distance in the lineage (e.g., B cells scored ∼0 on the Ery cell type score). All *in-vitro* samples had high Ery cell type scores --, yet lower than that of Ery samples in the reference. They also scored high for both MEP and CMP modules.

For deeper insight on the specificity of the effects of treatments, we juxtaposed pairs of modules. We observed that samples receiving HYDRO in round 1 had an increased Ery module score but left the MEP score unaffected (**Fig. 3e**). On the other hand, treatment with INS+HEP in the second round decreased the MEP module score – a desired effect – yet this came at the cost of a reduced Ery score (**Fig. 3f**) and an increased B cell module score (**Fig. 3g**).

Altogether, these observations link the previously observed effects of different treatments to concrete changes in the accessibility of specific chromatin regions and show that the quantification chromatin module scores provide a measure of the mixed identity of cells as a function of other cell types. The Ery module of cells generated by *in-vitro* differentiation is not as strongly accessible as in reference erythroblasts and coexists with residual progenitor chromatin programs, which hold them from fully achieving the erythroid cell state.

### Transcription factor analysis prioritises candidate culture components to correct the discrepancies between in-vitro and in-vivo cells

As established in the previous sections, acquisition of a full erythroblast identity consists of both opening of Ery-specific chromatin and closure of non-Ery chromatin. While our samples progressed towards such erythroblast identity (cp. **Fig. 1**), they did not fully acquire the erythroblast chromatin state even with the best-performing treatment (5E) (cp. **Fig. 3d-f**). A close inspection of the accessibility of chromatin modules after differentiation confirmed that many Ery-specific regions had gained accessibility, while many HSC-, CMP-, and lymphocyte-specific regions had lost accessibility (**Supplementary Fig. S8a**). However, regions in the MEP module and other non-Ery regions (AUC ≤ 0.05) remained accessible at the end of the differentiation protocol. We summarized these sets of regions as “concordant open”, “concordant closed”, and “discordant open”, respectively (**Fig. 4a**). The culture ingredients tested as part of our treatments did not sufficiently address the discordant regions, and we hypothesized that rectifying these discrepancies to the *in-vivo* target cells would likely improve fidelity of differentiation.

**Figure 4.**
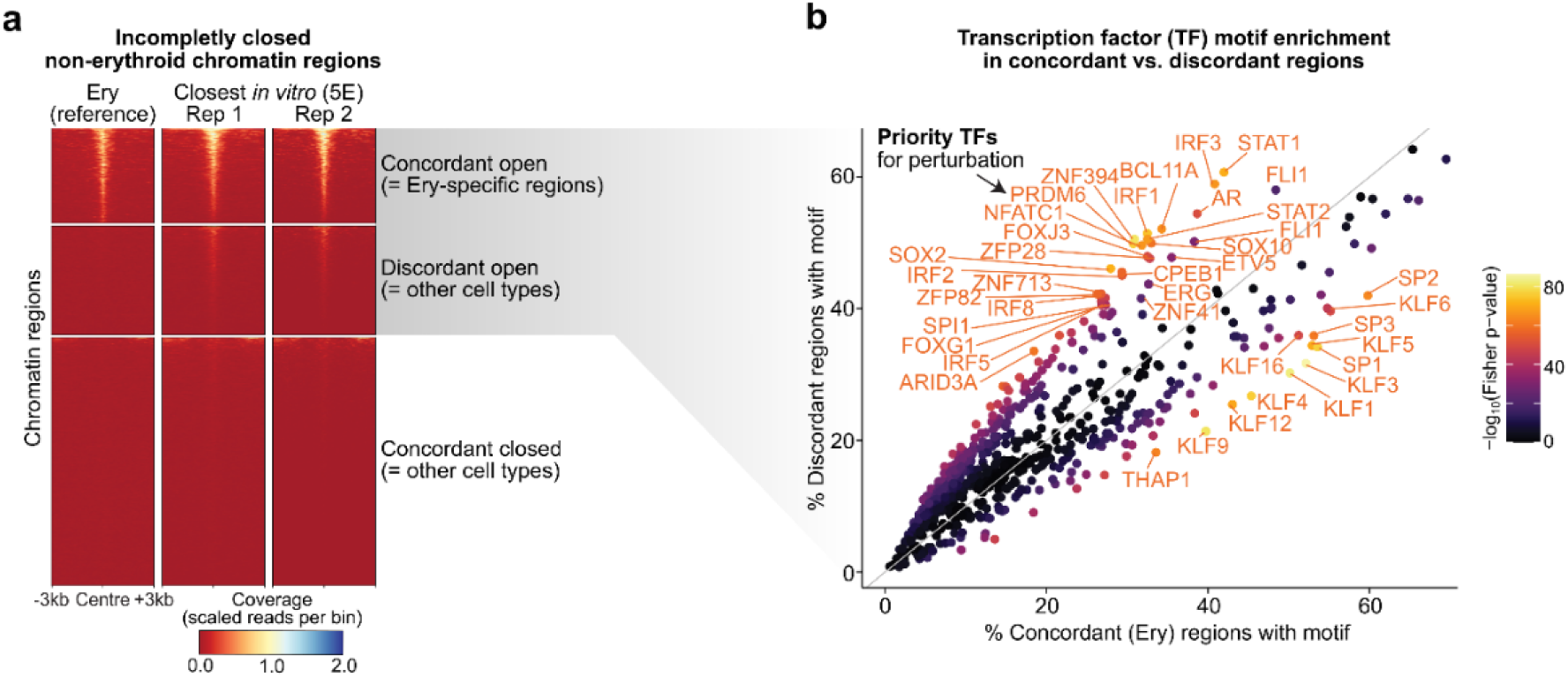
Mismatches persist in chromatin adaptation to erythroblasts even for best-performing protocols. **a**, Sorted region heatmaps comparing accessibility between erythroblast samples (Ery; one representative sample from (32)) and two replicate samples after differentiation with the best available treatment *in vitro* (treatment 5E = high erythropoietin over two rounds). Each block corresponds to cell type-specific chromatin modules that have been arranged by likeness to the Ery reference into concordantly open regions (top; identical to Ery-specific chromatin module), discordantly open (middle; MEP- and Non-Ery regions), and concordantly closed (bottom; HSPC and lymphocyte-specific regions), see Supplementary Figure S8a. Concordant regions (top and bottom) match between Ery and 5E, whereas discordant regions (middle) remain erroneously accessible in treatment 5E. **b**, Direct comparison of the percentage of regions with >0 transcription factor (TF) motifs from the HOCOMOCO (46) v11 database in the concordant (x axis) versus the discordant (y axis) regions. Colour represents the log10 p-value of a two-sided Fisher test calculated on a 2-dimensional contingency table for that motif (number of motif hits in 4, 872 discordant regions versus 4, 218 concordant regions). We label TFs with motifs absolute percentage changes ≥ 0.15, or those uniquely enriched in the discordant regions (percentage ≥ 40%) which were used for the enrichments below. The motif frequencies in concordant and discordant regions are provided in Supplementary Table S7.

We reasoned that targeting TFs that bind to the discordantly open regions was a promising strategy to rationally enhance treatment protocols. We therefore searched for DNA binding motifs that occurred more frequently in discordant regions compared to Ery regions (**Fig. 4b**). In agreement with our earlier analyses (cp. **Fig. 3c**), KLF motifs were abundant in the Ery regions and less pervasive in discordant regions (**Fig. 4b**). In contrast, we found motifs of interferon response factors (IRF1/2/3/5/8), STAT transcription factors (STAT1/2), and key regulators of HSPCs (ERG, FLI1) enriched in discordant regions (**Supplementary Table S7**).

Many of these TFs are regulators of other (non-erythroid) haematopoietic lineages (**Supplementary Fig. S8b-c**). We ranked TFs by the enrichment of their motifs and kept only those TFs with motifs in at least 40% of discordant regions as “priority TFs” (**Fig. 4b**, top region). No single motif was found in more 60% of the regions, suggesting that the discordant regions might be a collection of different functional programmes.

Next, we searched for ligand perturbations that could inhibit the expression of the selected TFs. To do this, we performed gene set enrichment analysis of the priority TFs against ligand perturbation databases using hypeR (57, 58) (**Fig. 5a, Supplementary Table S8**). Among the top hits resulting in a down-regulation of the TFs in other cell types, were multiple interleukins (TGF-β, IL-4, IL-10, IL-15) and steroid hormones (estradiol, testosterone) previously used by others for haematopoietic differentiation *in vitro* (59–66). These observations support the relevance of our selection. To validate the computational predictions, we tested the inclusion of TGF-β1 and IL-4 on top of the best-performing treatment 5E. Previous research showed that TGF-β impairs the proliferation of stem and erythroid cell cultures (61). To mitigate this, we opted for a low-dose and treated cells with 0.1 ng/ml TGF-β from day 1 or day 7. We analysed the conventional erythroid-associated markers CD71, CD36, and CD235a by flow cytometry to evaluate the degree of erythroid differentiation at day 7 (round 1) and day 11 (round 2) (**Fig. 5c-d, Supplementary Fig. S9a-c**). Already at day 7, we observed that the addition of 0.1 ng/ml TGF-β to the previously top-performing protocol led to a significant increase of CD71^+^/CD235a^+^ cells (immature erythroblasts) and a reduction of CD71^-^/CD36^+^ cells compared to the control set (**Fig. 5b, Supplementary Fig. S9b**). The addition of IL-4 also modestly enhanced the proportion of CD71^+^/CD36^+^ double-positive erythroid progenitors. However, the effect of IL-4 on erythropoiesis was not evident anymore after round 2, when assessing the marker expression by flow cytometry (**Fig. 5c, Supplementary Fig. S9c**). In contrast, the effects of TGF-β were even more pronounced at the end of the second round of differentiation (after 11 days), as shown by a significantly higher proportion of cells in an advanced maturation stage (CD71^+^/CD235a^+^). These effects were even stronger if TGF-β was administered from the beginning of differentiation (**Fig. 5d**), showing that differentiating cells had already advanced further on the erythropoietic trajectory at day 7 and consolidated in the CD71^+^/CD235a^+^ state at day 11 (**Supplementary Fig. S9d**). The improved differentiation outcome was not accompanied by a measurable loss in yield under using the low doses of TGF-β applied in our experiments (**Supplementary Fig. S9e**). To functionally test the effect of TGF-β and IL-4 on erythroid maturation, we measured the levels of *HBB* by RT-qPCR and observed that TGF-β increased *HBB* levels at day 11 and day 14 of differentiation (**Fig. 5e**). In agreement, pellets form TGF-β treatment appeared qualitatively redder, suggesting a higher haemoglobin content (**Supplementary Fig. S10a**). We then used Propidium Iodide (PI) and Hoechst staining to quantify the fraction of enucleated cells (indicating reticulocytes) and observed a mild, yet significant increase in double-negative cells at day 14 upon TGF-β treatment (**Fig. 5f, Supplementary Fig. S10b**); similarly, we observed a higher proportion of CD235a^+^ cells (**Supplementary Fig. S10c**). Finally, we performed Wright-Giemsa staining to measure cell morphology (**Fig. 5g**). In agreement with our flow cytometry analyses, cytospins showed a higher proportion of orthochromatic and polychromatic cells (late-stage erythroblasts) when cells were treated with TGF-β (**Fig. 5h**). These functional assays support the notion that our data-driven approach resulted in functionally more mature, differentiated cells.

**Figure 5.**
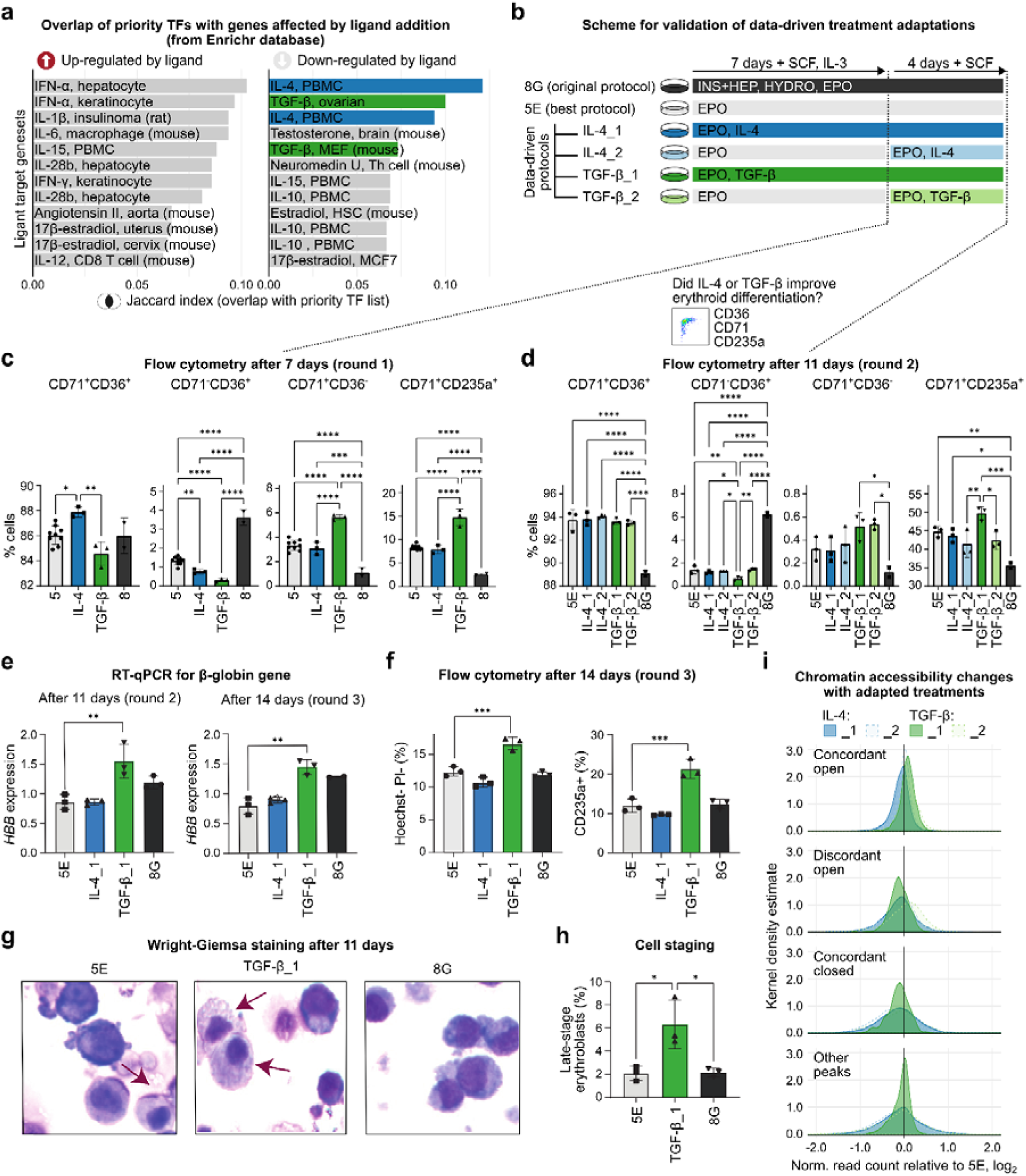
Data-driven selection of protocol modifications to correct chromatin mismatches between *in-vitro* and *in-vivo* cells. **a**, Gene set overlap analysis comparing TFs associated with the discordant-enriched motifs in **Fig. 4b** with genes affected by perturbations ligand perturbations from the Enrichr database (collections: “Ligand_Perturbations_from_GEO_down/up”) using hypeR (background: all 676 motifs in the HOCOMOCO database; all tested gene sets were restricted to genes in this background). Gene sets were ranked by the Jaccard index. Overlap results are provided in Supplementary Table S8. **b**, Schematic overview of validation experiments for data-driven improvements to Ery differentiation based on the enrichment results in panel (a). We tested the original protocol (8G), the best-performing protocol (5E), in comparison to a modified version of protocol 5E with the addition of IL-4 or TGF-ß in both rounds of differentiation (IL-4_1, TGF-ß_1) or only in round 2 (IL-4_2, TGF-ß_2). **c, d**, Flow cytometry results comparing Ery markers at round 1 (day 7, panel c) and round 2 (day 11, panel d) of differentiation. Each panel shows, in order: the percentage of CD71/CD36-double-positive cells, of CD36-positive cells, of CD71-positive cells, and of CD71/CD235a-double-positive cells. Statistical analyses using ordinary one-way ANOVA. *, p < 0.05; **, p < 0.01; ***, p < 0.001; ****, p < 0.0001. **e**, RT-qPCR analysis of *HBB* expression at day 11 and 14. Bars represent means and standard deviations of three biological replicates. Statistical analyses using ordinary one-way ANOVA. **, p < 0.01. **f**, Quantification of enucleated cells as Hoechst/PI double-negative cells by flow cytometry. Bars represent means and standard deviations of three biological replicates. Statistical analyses using ordinary one-way ANOVA. ***, p < 0.001. **g**, Wright-Giemsa of cytospin preparations of cells at day 11 of erythroid differentiation. Late erythroblast cells are marked with an arrow. **h**, Quantification of late erythroblast cells upon treatment with TGF-ß or IL-4. Bars represent means and standard deviation of three biological replicates. One thousand cells were counted and staged per replicate. Statistical analyses using ordinary one-way ANOVA. *, p < 0.05. **i**, Distribution of log_2_ fold changes per peak group (compared to all other peaks) in each adapted protocol compared to treatment 5E. Green and blue tones correspond to TGF-β and IL-4 treatment, respectively. Filled and dotted lines correspond to the treatment applied in both rounds (TGF-β_1, IL-4_1), or only at round 2 (TGF-β_2, IL-4_2), respectively.

To assess whether the observed advancement of erythropoiesis was linked to changes at chromatin level, we performed ATAC-seq of at the end of round 2 of the modified differentiation protocols in comparison to treatment 5E (data overview and quality metrics in **Supplementary Table S1**). We found that chromatin accessibility was globally very similar to the base protocol 5E (**Supplementary Fig. S10d**). In comparison to 5E, TGF-β supplementation throughout differentiation (*TGF-*β*_1*) yielded a further increase in the accessibility of concordantly open and a further decrease of accessibility in concordantly closed regions, respectively (**Fig. 5i**). Regions that previously retained accessibility (discordantly open), were less accessible. However, these changes were small, indicating that only a fraction of cells was affected (e.g., median log_2_ fold change TGF-β_1 vs 5E in discordant peaks = −0.2). Only 42 peaks were significantly affected in at least one of the adapted treatments (DESeq2 (42), FDR-adjusted P-value ≤ 0.05; **Supplementary Table S9**). While most of these peaks had matches to motifs of priority TFs (**Supplementary Fig. S10e**), it is unclear from our data whether this had a functional relevance since the same motifs are common among peaks that are not significantly affected (FDR-adjusted P-value > 0.05). Rather, the primary mode of action of the TGF-β (which is not a TF or epigenetic modifier) on erythropoiesis might be not via directly altering chromatin but by modulation of the activities of TFs binding in those regions. Changes in chromatin would likely be a secondary effect resulting from advanced differentiation leading the progressive closing down of regions, like for instance for the promoter of *Rho GTPase activating protein 10* (*ARHGAP10*), which is gradually closing throughout erythropoiesis *in vivo* and which is among the significantly affected discordant peaks in our data (**Supplementary Fig. S10f**).

Altogether, our results confirm that supplementation of our tested differentiation protocols with TGF-β improves erythroblast production. This example showcases how in-depth analysis of chromatin during *in-vitro* differentiation not only allows to rank treatments by similarity to the reference, but can also provide a rational, data-driven shortlist of additional treatments tailored to overcome shortcomings in chromatin programming in differentiation protocols.

## DISCUSSION

In this study, we introduced an approach for optimizing directed differentiation of stem cells by iteratively approximating the chromatin state of target cells. Applied to the differentiation of haematopoietic stem cells (HSCs) into erythroblasts, we ranked differentiating cells using chromatin distance as a simple, objective measure of similarity. Our results indicate that a greedy strategy, which iteratively selects the treatment closest to the target at each step, yielded the optimal protocol, while simplifying the experimental design and reducing workload. The consistency between chromatin distance and conventional cell fidelity markers (CD36, CD71) at the end of differentiation validates this approach.

Moreover, with access to a full chromatin accessibility profile, we exposed residual discrepancies to the desired target cells even in the best-performing protocols. The observed deviations might indicate meta-stable intermediate cell states in which competing lineage programmes have not yet been inactivated completely (67–69). Lacking continuous interactions with the surrounding niche to refine and stabilize signalling, the precision of differentiation trajectories diminishes *in vitro*. Additional rounds of differentiation or alternative treatments may be necessary to close this gap. Our analysis identified specific chromatin regions associated with myeloid progenitors that were incompletely closed in differentiated cells. Harnessing discrepancies in the footprints of specific transcription factors (e.g., STAT1/2 and IRF1/2/8), allowed evidence-based prioritization of candidate perturbations that may help to achieve the desired erythroid cell state, amongst them IL-4 and TGF-β. Consistent with these bioinformatic predictions, low doses of TGF-β have previously been reported to promote erythropoiesis (60). In contrast, it was also shown that high concentrations of TGF-β might downregulate erythrocyte production by inhibiting erythroblast proliferation (61). Our validation experiments proved that supplementation of the best-performing protocol 5E (SCF, 3IU EPO) with low-dose TGF-β (0.1 ng/ml), especially if administered from the beginning of differentiation, fast-tracked erythroid differentiation with a significantly increased yield of CD71^+^/ CD235a^+^ cells already after 11 days of differentiation. This confirms the validity of our approach for data-driven prioritization of modifications to existing differentiation protocols.

Our work demonstrates the potential of chromatin-guided, greedy selection for simplifying and enhancing differentiation protocols. The methodology can be applied even at low sequencing depth lending itself to high multiplexing (Supplementary Fig. S4). We found chromatin distance anti-correlated with proximity to target cell type in multiple, but not all haemopoietic cell types (Supplementary Fig. S5), arguing the approach may be extended to other cell types that are challenging to produce *in vitro*, if appropriate care is taken. Moreover, the approach can conceptually also be extended to transdifferentiation and reprogramming – although further work is required to confirm validity in these settings. The main challenge to be solved for any new target cell type is to define the target chromatin state and a “neighbourhood” of alternative (undesirable) cell fates, as a reference for chromatin distance. With the increasing availability of multi-modal single-cell reference atlases, defining such cell states will soon be feasible even for cell types that were previously poorly described and without existing differentiation protocols (34, 37, 70–73). A selection of treatment options (including culture components as tested here, but perhaps also including other chemical and mechanical stimulants, co-cultures, etc.) may be selected from prior knowledge or using data-driven selection between starting and target cells, as we have done here in the final steps of our protocols. The computational pipeline we developed facilitates these analyses and comparison of test samples to defined target cell types.

Despite the advantages presented here, our approach has limitations: first, we cannot exclude that the greedy selection may in some cases result in solutions which are not the global optimum. We have chosen a chromatin-based distance as an objective function because it captures the genome-wide meta-stable epigenetic state of cells and deemphasises temporary fluctuations (compared to mRNA expression). While prior research argues for the superiority of chromatin accessibility measurements over mRNA expression for distinguishing haematopoietic cell types (33), we have not demonstrated this superiority in our experiments. However, no one-dimensional function can weigh all aspects of the underlying biology thoroughly; hence, complementary functional assays are important to evaluate the final cell products. Relatedly, we defined chromatin distance as a linear metric in Euclidean PCA space. We are aware that this is an oversimplification. *In vivo*, differentiation often proceeds via multiple progenitor stages that may not immediately resemble the final target cell. In such cases, chromatin distance can be used stepwise to approximate intermediates first before proceeding to the next differentiation process. In future, more complex, non-linear chromatin distance models could help directly factorize differentiation stages and technical covariates into a compound distance metric. Second, we did not directly address the heterogeneity of differentiated cells but rather opted for a bulk assay as a straightforward, cost-effective solution. This does not impair the search for homogeneous cell products, since the average genome-wide profiles of mixed cell populations will deviate from the specific target profile, which would lead to protocols resulting in mixed populations being deselected by our greedy optimization strategy. In practice, we recommend pairing ATAC-seq with flow cytometric or imaging-based assays at single-cell resolution to confirm homogeneity of differentiated cells. Finally, scalability and practicability of the generation of products for cell therapies depends on technical factors beyond the phenotypic approximation.

As a proof-of-principle, we have decided to work here with an exhaustive setup testing all combinations of 8×8 treatments in two rounds of sequential treatments, and to trace back the data generated during the process to retrospectively evaluate what result the greedy selection would have yielded. In an actual application, this optimization may be done in cycles for an arbitrary number of treatment rounds, evaluating the result after round 1 first, restarting with the best treatment and varying treatments at round 2, again picking the best solution, and so on. This works well with a limited number of rounds but remains a time-consuming procedure. Conceptually, treatment evaluation and readjustment may be done live throughout an ongoing experiment: our computational simulations (Supplementary Fig. S4) suggest that even low-coverage ATAC-seq (e.g., with current benchtop sequencing machines) would be sufficient to estimate chromatin distance, potentially allowing best-treatment selection in between steps of an ongoing differentiation protocol (typically 2-5 days). However, while this methodology allows prioritization of candidate treatments, chromatin distance does not necessarily translate directly into physiological relevance. It is therefore crucial to test all prioritized protocols by validating protein markers and by examining functional characteristics of the generated cell products.

## DATA AVAILABILITY

ATAC-seq data related to this project was deposited at the Gene Expression Omnibus (GEO <u>GSE291386</u>). Genomic data from this study can be accessed interactively via the UCSC genome browser at https://genome.ucsc.edu/cgi-bin/hgTracks?hubUrl=http://fh.ccri.at/ML2Cell_trackhub/hub.txt, or using a pre-configured IGV session at https://fh.ccri.at/ML2Cell_trackhub/igv_session_complete.xml.

## ACKNOWLEDGEMENTS

We thank Eva Bozsaky and Paul Batty for their support with microscopy.

## MATERIAL & METHODS

### Human erythroid cell culture

G-CSF-mobilized human peripheral blood CD34+ cells (Stem Cell Technologies) were differentiated into immature erythroid cells utilizing a three-phase culture protocol. For standardisation, we used the same batch of cells for all experiments described in this study. Briefly, cells were thawed and cultured overnight at a density of 10^4^ cells/ml in X-VIVO 10 serum-free haematopoietic cell medium (Lonza) supplemented with 100 ng/ml recombinant human SCF (Biotechne), 100 ng/ml recombinant human TPO and 50 ng/ml recombinant human FLT3 Ligand (PeproTech). The next day, cells were washed and resuspended at a density of 1-2.5 × 10^4^ cells/ml into erythroid differentiation medium 1 (EDM1) consisting of IMDM supplemented with 100 ng/ml hSCF, 330 μg/ml human holo-Transferrin, 5 ng/ml recombinant human IL-3 (PeproTech), 1% penicillin/streptomycin, 5% heat-inactivated human serum (Sigma) and 1 or 3 IU human recombinant EPO (Stem Cell Technologies). When it is indicated, 1 μM hydrocortisone, 10 μg/ml recombinant human insulin and 2 IU/ml heparin (Sigma) were added as well. From day 7 to day 11, cells were cultured at a concentration of 10^5^ cells/ml in EDM2 (EDM1 without hIL-3). From day 11 to day 14, cells were cultured in EDM3 (EDM2 without hSCF) at a concentration of 0.75-1.5 × 10^6^ cells/ml in the presence of 1 mg/ml Holo-Transferrin. For validation experiments, CD34+ cells were differentiated into immature erythroid cells as described above, with the addition of 0.1 ng/ml human TGF-β1 recombinant protein (PeproTech) or 10 ng/ml human recombinant IL-4 (PeproTech) in EDM1 and EMD2 when indicated. Cells were cultured at 37°C and 5% CO2. These materials are listed in **Supplementary Table S10**.

### Flow Cytometry

Cells were washed in PBS supplemented with 1% FBS and incubated with antibodies for 30 minutes at 4°C. The following antibodies were used: FITC-conjugated CD71 antibody (clone OKT9, eBioscience, 1:40 dilution); APC-conjugated CD36 antibody (clone 5-271, Biolegend, 1:40 dilution) and BV421-conjugated CD235a antibody (clone HIR2 GA-R2, BD, 1:20 dilution). Cells were stained using 1 µg/ml Hoechst 33342 (#H3570, Thermo) for 20 minutes at 37 °C, followed by staining with 4 µg/ml Propodium Iodide (#81845, Sigma) for 5 min at 4 °C to assess enucleation. Flow cytometry was conducted on a BD Bioscience Fortessa instrument and analysed using FlowJo software v10.10.

### RNA isolation and RT-qPCR

RNA was harvested from 1-2 million cells using the RNeasy Plus Mini Kit (Qiagen, #74134). 500 ng of RNA was used for cDNA synthesis using the iScript cDNA Synthesis Kit (BioRad, #1708891). qPCR was performed using the iQ5 SYBR Green Supermix (BioRad, #1708885) and ran in the CFX Connect Real Time System (BioRad). An initial denaturation at 95 °C for 3 min, followed by 40 cycles of 10 s denaturation at 95 °C and 30 s annealing and extension at 60 °C was performed using the following oligonucleotides for *HBB* (CTGAGGAGAAGTCTGCCGTTA; AGCATCAGGAGTGGACAGAT) and *RPLP0* (GATTGGCTACCCAACTGTTG; CAGGGGCAGCAGCCACAAA).

### Wright-Giemsa staining

Approximately 0.1 million cells were immobilized onto microscope slides using a low-speed centrifuge (Hettich Rotofix) and a funnel at 260 rpm for 8 min. Next, the funnel and media were removed, and slides were spun for another 4 min at 330 rpm. The cells were then fixed in methanol for 1 min, immerged in modified Wright-Giemsa (WG16, Sigma) for 2 min, followed by incubation in phosphate buffer (pH 6.8; #1113740100, Merck) for 4 min. Finally, slides were rinsed in ddH20 and air-dried. Slides were imaged using a ZEISS Axio Imager Z2 using 20x and 40x objectives. One thousand cells for each slide were manually counted and scored according to morphology.

### Chromatin accessibility analysis (ATAC-seq)

#### Library preparation and sequencing

For ATAC-seq, 50, 000 cells were collected at day 7 and day 11, in biological duplicates. Cells were washed using cold PBS and subsequently lysed in cold lysis buffer (10 mM Tris-HCl pH 7.4, 10 mM NaCl, 3 mM MgCl2, 0.1% IGEPAL CA-360). Immediately after lysis, nuclei were collected by centrifugation, and the pellet was resuspended in 50 μl transposase reaction mix (Illumina Nextera DNA Preparation kit). The transposition reaction was carried out at 37°C for 30 minutes. Following transposition, the samples were purified and eluted in 10 μl Elution Buffer (Qiagen Minelute PCR purification kit). Libraries were amplified by PCR for a minimum of 5 cycles. Additional cycles were added to reach exponential amplification according to qPCR. Oligonucleotides are provided in **Supplementary Table S11**. Libraries were purified and quantified using a Qubit dsDNA HS Assay kit (Invitrogen). Before pooling, the quality of the library was determined by fragment analysis on a TapeStation 4200 (Agilent Technologies) using a High Sensitivity D1000 ScreenTape. Samples were sequenced on a NovaSeq X Plus to obtain at least 30M PE150 reads per library. **Supplementary Table S1** provides an overview of all sequence data and associated quality metrics.

#### Data preprocessing and quality control

Bioinformatic pipelines were obtained using nextflow (v22.10.6) nf-core (v2.14.1) (41). Raw chromatin accessibility data from purified cell types in the bone marrow was obtained from a previously published work (32), GEO GSE74912) using the nexftlow (v23.10.1) nf-core pipeline nf-core/fetchngs (v1.10.0). For alignment, peak calling, quality control and accessibility count matrices, the reference and test datasets were processed separately using the nf-core/atacseq pipeline (v2.1.2) (directives: --broadpeak, --aligner bwa, macs_genome_size: 2749859687, iGenomes hg38). To obtain accessibility counts in the reference regions for both datasets, test reads were re-counted on the reference consensus peak set using a custom workflow employing the nf-core module subread/featureCounts. Samples with a Fraction of reads in peaks (FRiP) score ≥ 0.05 were kept for downstream analysis. The coordinates and annotations of peaks used in this study are provided in **Supplementary Table S4**.

#### Normalisation and selection of cell type-informative PCA components and calculation of distances to target

All subsequent analyses were implemented in R (v4.1.3) and Bioconductor (v3.14). Depth-normalised counts on 595, 523 regions for all samples were obtained for one reference dataset and two test datasets using DEseq2 (42) (v1.34.0).

Normalised counts for the reference cell types were subjected to principal component analysis. Loadings for the reference samples on 80 principal components were then used to project the test samples onto this PCA space. Reference sample PCA embeddings were used to train a Random Forest multiple classification model using the R library RandomForest (v4.7-1, ntrees=500). Classification error estimations, variable importance, and cross-validation were performed by performing out-of-bag (OOB) analysis while building the decision trees. Components with a mean decrease in Gini index of ≥ 2 (high-importance PCs) were kept for a second round of Random Forest to quantify the decrease in classification error. High-importance PCs were used to calculate the Euclidean distance between each sample and the centroid (multidimensional mean) of erythroblasts.

#### Computational down-sampling analysis

To simulate the effects of varying sequencing depth in the test and reference dataset, we calculated the average sequencing depth of the test dataset (our ATAC-seq data) and the reference haematopoietic dataset and randomly down-sampled read counts assigned to each peak to defined read depths (0.5, 1, 5, 10, 15, 20, 25 million reads for the test data; 1, 10, 20 million reads for the reference) as a fraction of the original read depth. We performed 250 random trials and evaluated the chromatin distances to the erythroid sample in all pairwise comparisons of test and reference read depth after selecting informative PCs as described above.

#### Evaluation of chromatin distance in other haematopoietic lineages

Sequencing read datasets comprising *in-vitro* samples from the respective publications (43–45) were downloaded and processed using the nextflow nfcore/atacseq pipeline, and subsequently reads in the reference peaks were counted using our custom nextflow workflow with the same parameters as described above. Instead of projecting each dataset to the reference PCA, a joint PCA with all the datasets was performed, which mitigated issues due to dataset quality control differences. For the calculation of the chromatin distance, high importance PCs (see above) with mean decrease Gini index above 1.2 were used (PC2, PC13, PC7, PC4, PC16, PC10, PC17, PC5, PC6, PC14, PC11, PC8, PC3, PC9 und PC12) to calculate the Euclidean distance between each sample and the PCA centroid of the samples the target cell type.

#### Selection of chromatin cell identity modules using the Receiver-Operating characteristic (ROC)

Chromatin regions with the highest loadings were collected from the first 10 PCs of the dataset and the high-importance PCs (see previous section). For each PC, regions were ranked a lower limit in the loading was set by a heuristic elbow metric (**Supplementary Fig. S7a**). Within this subset, up to 8000 peaks per PC were scrutinised exhaustively for cell type signals using ROC analysis. ROC curves and their respective area under the curve (AUC) were then calculated for each high-loading region and cell type using the normalised accessibility counts using the R package pROC (v1.18.0). Regions with an AUC ≥ 0.95 are used as cell identity chromatin modules (**Fig. 3**) and regions with AUC ≤ 0.05 are also kept as distinctively closed for that cell type. Test samples from *in-vitro* differentiation were left out of the ROC computation. AUC values for all peaks are provided in **Supplementary Table S4**.

#### Mixed-effects modelling of treatment effects

Mixed-effects modelling was performed in R using the function *lmer* from the package lme4 (v1.1-29). The model evaluated the contributions of fixed effect predictors (1.EPO and 2.EPO) and random effect predictors (1.HYDRO, 2.HYDRO, 1.INS+HEP, 2.INS+HEP) on the response (the distance to the Ery centroid). The intercepts and coefficients of the model are provided in **Supplementary Table S3**.

#### Transcription factor binding motif counting and enrichment analysis

Transcription factor motifs in all regions were counted R package motifmatchr (v1.16.0) using position-specific scoring matrices from the HOCOMOCO database (46) (v11). A region with >0 motif hits for a TF was considered positive. Then the number of positive regions in a selected groups was compared to the background in question. In the case of transcription factors enriched in cell-type specific regions, the background was all the regions. In the case of the discordant vs concordant regions, the background was the concordant regions. Fold-enrichments were calculated using the formula log2(number of positive regions in sample/ number of positive regions in background). Statistical assessment of the enrichments was performed via a right tailed hypergeometric test (finding the probability of sampling X or more positive regions without replacement in a sample of n regions given an expected number of p(n) positive regions) and a Fisher test on the contingency matrix. Probabilities were then adjusted for multiple testing using the Benjamini-Hochberg procedure. Motif enrichment results are provided in **Supplementary Tables S6 and S7**.

#### Chromatin cell identity module scoring and analysis

Single sample gene set variation analysis scoring was performed using the method ‘ssgsva’ in from the R package GSVA (47) (v1.42.0). For each cell type, cell identity regions were treated as genes for this analysis and scores were calculated from the normalised count matrix for all samples including the test (**Fig. 3e**). To obtain a relative score (**Fig. 3d**), the average score for hematopoietic stem cell samples was subtracted and the value was divided by the average score of the query cell type. For example, to calculate the B cell module score for an Ery sample, the average B cell score for the HSCs was first subtracted from all samples and then divided by the B cell score of B cells. Negative scores were set to zero. The HSC module score is not defined for all samples using this formula.

#### Gene set enrichment analysis of closest transcription start sites

The closest transcription start site for the regions in the analysis was found using the GenomicRanges package (v1.46.1), specifically the function distanceToNearest (ignore.strand=T). The resulting genes per group were submitted to a gene set enrichment analysis using the package hypeR (48) (v 1.10.0, test=”hypergeometric”) versus different gene sets and were adjusted for multiple testing using the Benjamini-Hochberg procedure, then filtered on p value (p ≤ 0.05) and sorted by overlap using the Jaccard index (size of the gene list intersection/size of the gene list union). Gene set enrichment results are provided in **Supplementary Tables S5 and S8**.

#### Generation of sorted region heatmaps

Sorted region heatmaps were generated with the Deeptools package (v3.5.4), specifically the tools computeMatrix (reference-point, -a=3000, b=3000, sortRegions, descend=S, bs=25, missingDataAsZero, referencePoint=center, p=10) and plotHeatmap (zMin=0, zMax=2, yMin=0, yMax=1). The reference sample for sorting was always the leftmost one in the plots.

#### Concordant and discordant region analysis

This analysis compared the opening and closing of peaks in the best protocol samples (those of 5E) relative to a true erythroblast sample (SRX1427827), as shown in **Figure 5a**. To separate concordant and discordant regions, all regions that could be informative of cell fidelity were used. This involved cell identity regions for all cell types (ROC AUC ≥ 0.95) and the negative cell identity regions for Ery (ROC AUC for Ery ≤ 0.05). To simplify clustering (which performed poorly with all the peaks together), blocks of peaks were visually split into mostly concordant open (all Ery peaks), concordant closed and mostly discordant (Ery negative peaks plus MEP peaks). Then, to further refine the discordant peak block, discordant regions were re-sorted and clustered using DeepTools plotHeatmap (hclust=4) using the aforementioned Ery sample as a sorting and clustering reference. This yielded three small clusters with concordant patterns and one large discordant cluster, which was then used as the discordant peaks. Similar treatment of the concordant or discordant groups yielded no obvious improvement on the group quality and thus both groups were kept according to the visual split.

#### Analysis of chromatin changes in adapted differentiation protocols

To compare the adapted protocols (TGF-β_1, TGF-β_2, IL-4_1, IL-4_2) versus the previously winning protocol (5E) we used DESeq2 (v1.34.0). Each condition was compared individually versus the samples of 5E. To call differentially accessible peaks, we tested only concordance peaks (concordant open/closed and discordant) with 5 or more reads in at least two samples of at least one condition. Peaks with an adjusted P-value less than or equal to 0.05 (using Benjamini-Hochberg correction) were reported as significant (**Supplementary Table S9**).

## REFERENCES

1. Mao, J., Saiding, Q., Shutong Qian, |, Liu, Z., Zhao, B., Zhao, Q., Lu, B., Mao, X., Zhang, L., Zhang, Y., et al. (2022) Reprogramming stem cells in regenerative medicine. Smart Medicine, 1, e20220005.

2. Cohen, D.E. and Melton, D. (2011) Turning straw into gold: directing cell fate for regenerative medicine. Nature Reviews Genetics 2011 12:4, 12, 243–252.

3. Cahan, P., Li, H., Morris, S.A., Lummertz Da Rocha, E., Daley, G.Q. and Collins, J.J. (2014) CellNet: Network biology applied to stem cell engineering. Cell, 158, 903–915.

4. Kirouac, D.C. and Zandstra, P.W. (2008) The Systematic Production of Cells for Cell Therapies. Cell Stem Cell, 3, 369–381.

5. Murry, C.E. and Keller, G. (2008) Differentiation of Embryonic Stem Cells to Clinically Relevant Populations: Lessons from Embryonic Development. Cell, 132, 661–680.

6. Liu, S., Striebel, J., Pasquini, G., Ng, A.H.M., Khoshakhlagh, P., Church, G.M. and Busskamp, V. (2021) Neuronal Cell-type Engineering by Transcriptional Activation. Front Genome Ed, 3, 715697.

7. Morris, S.A., Cahan, P., Li, H., Zhao, A.M., San Roman, A.K., Shivdasani, R.A., Collins, J.J. and Daley, G.Q. (2014) Dissecting engineered cell types and enhancing cell fate conversion via Cellnet. Cell, 10.1016/j.cell.2014.07.021.

8. Poldee, S., Metheetrairut, C., Nugoolsuksiri, S., Frayne, J. and Trakarnsanga, K. (2018) Optimization of an erythroid culture system to reduce the cost of in vitro production of red blood cells. MethodsX, 5, 1626–1632.

9. Toms, D., Deardon, R. and Ungrin, M. (2017) Climbing the mountain: Experimental design for the efficient optimization of stem cell bioprocessing. J Biol Eng, 11, 1–10.

10. Yeo, G.H.T., Lin, L., Qi, C.Y., Cha, M., Gifford, D.K. and Sherwood, R.I. (2020) A Multiplexed Barcodelet Single-Cell RNA-Seq Approach Elucidates Combinatorial Signaling Pathways that Drive ESC Differentiation. Cell Stem Cell, 26, 938–950.e6.

11. Tarunina, M., Hernandez, D., Johnson, C.J., Rybtsov, S., Ramathas, V., Jeyakumar, M., Watson, T., Hook, L., Medvinsky, A., Mason, C., et al. (2014) Directed Differentiation of Embryonic Stem Cells Using a Bead-Based Combinatorial Screening Method. PLoS One, 9, e104301.

12. Kim, H.D., Lee, E.A., Choi, Y.H., An, Y.H., Koh, R.H., Kim, S.L. and Hwang, N.S. (2016) High throughput approaches for controlled stem cell differentiation. Acta Biomater, 34, 21–29.

13. Smucker, B., Krzywinski, M. and Altman, N. (2018) Optimal experimental design. Nat Methods, 15, 559–560.

14. Rao, R.S., Kumar, C.G., Prakasham, R.S. and Hobbs, P.J. (2008) The Taguchi methodology as a statistical tool for biotechnological applications: A critical appraisal. Biotechnol J, 3, 510–523.

15. Rao, R.S., Kumar, C.G., Prakasham, R.S. and Hobbs, P.J. (2008) The Taguchi methodology as a statistical tool for biotechnological applications: A critical appraisal. Biotechnol J, 3, 510–523.

16. Yasui, R., Sekine, K., Yamaguchi, K., Furukawa, Y. and Taniguchi, H. (2021) Robust parameter design of human induced pluripotent stem cell differentiation protocols defines lineage-specific induction of anterior-posterior gut tube endodermal cells. Stem Cells, 39, 429–442.

17. Marinho, P.A., Chailangkarn, T. and Muotri, A.R. (2015) Systematic optimization of human pluripotent stem cells media using Design of Experiments. Scientific Reports 2015 5:1, 5, 1–13.

18. Yasui, R., Sekine, K. and Taniguchi, H. (2021) Clever Experimental Designs: Shortcuts for Better iPSC Differentiation. Cells 2021, Vol. 10, Page 3540, 10, 3540.

19. Li, L., Tan, D., Liu, S., Jiao, R., Yang, X., Li, F., Wu, H. and Huang, W. (2020) Optimization of Factor Combinations for Stem Cell Differentiations on a Design-of-Experiment Microfluidic Chip. Anal Chem, 92, 14228–14235.

20. García, A. (2024) Greedy algorithms: a review and open problems.

21. Farnsworth, S.L., Qiu, Z., Mishra, A. and Hornsby, P.J. (2019) Optimization of Differentiation of Nonhuman Primate Pluripotent Cells Using a Combinatorial Approach. Methods Mol Biol, 1919, 187–197.

22. Cahan, P., Cacchiarelli, D., Dunn, S.J., Hemberg, M., de Sousa Lopes, S.M.C., Morris, S.A., Rackham, O.J.L., del Sol, A. and Wells, C.A. (2021) Computational Stem Cell Biology: Open Questions and Guiding Principles. Cell Stem Cell, 28, 20–32.

23. Godoy, P., Schmidt-Heck, W., Hellwig, B., Nell, P., Feuerborn, D., Rahnenführer, J., Kattler, K., Walter, J., Blüthgen, N. and Hengstler, J.G. (2018) Assessment of stem cell differentiation based on genome-wide expression profiles. Philosophical Transactions of the Royal Society B: Biological Sciences, 373.

24. Ding, J., Alavi, A., Ebrahimkhani, M.R. and Bar-Joseph, Z. (2021) Computational tools for analyzing single-cell data in pluripotent cell differentiation studies. Cell reports methods, 1.

25. Kumar, P., Tan, Y. and Cahan, P. (2017) Understanding development and stem cells using single cell-based analyses of gene expression. Development, 144, 17–32.

26. Lo, E.K.W., Velazquez, J.J., Peng, D., Kwon, C., Ebrahimkhani, M.R. and Cahan, P. (2023) Platform-agnostic CellNet enables cross-study analysis of cell fate engineering protocols. Stem Cell Reports, 18, 1721–1742.

27. Kinney, M.A., Vo, L.T., Frame, J.M., Barragan, J., Conway, A.J., Li, S., Wong, K.-K., Collins, J.J., Cahan, P., North, T.E., et al. (2019) A systems biology pipeline identifies regulatory networks for stem cell engineering. Nat Biotechnol, 37, 810–818.

28. Loh, K.M., Ang, L.T., Zhang, J., Kumar, V., Ang, J., Auyeong, J.Q., Lee, K.L., Choo, S.H., Lim, C.Y.Y., Nichane, M., et al. (2014) Efficient endoderm induction from human pluripotent stem cells by logically directing signals controlling lineage bifurcations. Cell Stem Cell, 14, 237–252.

29. Kamimoto, K., Stringa, B., Hoffmann, C.M., Jindal, K., Solnica-Krezel, L. and Morris, S.A. (2023) Dissecting cell identity via network inference and in silico gene perturbation. Nature, 614, 742–751.

30. Kamimoto, K., Adil, M.T., Jindal, K., Hoffmann, C.M., Kong, W., Yang, X. and Morris, S.A. (2023) Gene regulatory network reconfiguration in direct lineage reprogramming. Stem Cell Reports, 18, 97–112.

31. Jung, S., Appleton, E., Ali, M., Church, G.M. and del Sol, A. (2021) A computer-guided design tool to increase the efficiency of cellular conversions. Nat Commun, 12, 1–12.

32. Corces, M.R., Buenrostro, J.D., Wu, B., Greenside, P.G., Chan, S.M., Koenig, J.L., Snyder, M.P., Pritchard, J.K., Kundaje, A., Greenleaf, W.J., et al. (2016) Lineage-specific and single-cell chromatin accessibility charts human hematopoiesis and leukemia evolution. Nature Genetics 2016 48:10, 48, 1193–1203.

33. Corces, M.R., Buenrostro, J.D., Wu, B., Greenside, P.G., Chan, S.M., Koenig, J.L., Snyder, M.P., Pritchard, J.K., Kundaje, A., Greenleaf, W.J., et al. (2016) Lineage-specific and single-cell chromatin accessibility charts human hematopoiesis and leukemia evolution. Nat Genet, 48, 1193–1203.

34. Zhang, K., Hocker, J.D., Miller, M., Hou, X., Chiou, J., Poirion, O.B., Qiu, Y., Li, Y.E., Gaulton, K.J., Wang, A., et al. (2021) A single-cell atlas of chromatin accessibility in the human genome. Cell, 184, 5985–6001.e19.

35. Buenrostro, J.D., Giresi, P.G., Zaba, L.C., Chang, H.Y. and Greenleaf, W.J. (2013) Transposition of native chromatin for fast and sensitive epigenomic profiling of open chromatin, DNA-binding proteins and nucleosome position. Nat Methods, 10, 1213–1218.

36. Dunham, I., Kundaje, A., Aldred, S.F., Collins, P.J., Davis, C.A., Doyle, F., Epstein, C.B., Frietze, S., Harrow, J., Kaul, R., et al. (2012) An integrated encyclopedia of DNA elements in the human genome. Nature, 489, 57–74.

37. Domcke, S., Hill, A.J., Daza, R.M., Cao, J., O’Day, D.R., Pliner, H.A., Aldinger, K.A., Pokholok, D., Zhang, F., Milbank, J.H., et al. (2020) A human cell atlas of fetal chromatin accessibility. Science (1979), 370, eaba7612.

38. Kiani, K., Sanford, E.M., Goyal, Y. and Raj, A. (2022) Changes in chromatin accessibility are not concordant with transcriptional changes for single-factor perturbations. Mol Syst Biol, 18, 10979.

39. Marstrand, T.T. and Storey, J.D. (2014) Identifying and mapping cell-type-specific chromatin programming of gene expression. Proc Natl Acad Sci U S A, 111, E645–E654.

40. Ma, S., Zhang, B., LaFave, L.M., Earl, A.S., Chiang, Z., Hu, Y., Ding, J., Brack, A., Kartha, V.K., Tay, T., et al. (2020) Chromatin Potential Identified by Shared Single-Cell Profiling of RNA and Chromatin. Cell, 183, 1103–1116.e20.

41. Ewels, P.A., Peltzer, A., Fillinger, S., Patel, H., Alneberg, J., Wilm, A., Garcia, M.U., Di Tommaso, P. and Nahnsen, S. (2020) The nf-core framework for community-curated bioinformatics pipelines. Nature Biotechnology 2020 38:3, 38, 276–278.

42. Love, M.I., Huber, W. and Anders, S. (2014) Moderated estimation of fold change and dispersion for RNA-seq data with DESeq2. Genome Biol, 15, 550.

43. Li, K., Wu, Y., Li, Y., Yu, Q., Tian, Z., Wei, H. and Qu, K. (2020) Landscape and Dynamics of the Transcriptional Regulatory Network During Natural Killer Cell Differentiation. Genomics Proteomics Bioinformatics, 18, 501–515.

44. Ramirez, R.N., El-Ali, N.C., Mager, M.A., Wyman, D., Conesa, A. and Mortazavi, A. (2017) Dynamic Gene Regulatory Networks of Human Myeloid Differentiation. Cell Syst, 4, 416–429.e3.

45. Ludwig, L.S., Lareau, C.A., Bao, E.L., Nandakumar, S.K., Muus, C., Ulirsch, J.C., Chowdhary, K., Buenrostro, J.D., Mohandas, N., An, X., et al. (2019) Transcriptional States and Chromatin Accessibility Underlying Human Erythropoiesis. Cell Rep, 27, 3228–3240.e7.

46. Kulakovskiy, I. V., Vorontsov, I.E., Yevshin, I.S., Sharipov, R.N., Fedorova, A.D., Rumynskiy, E.I., Medvedeva, Y.A., Magana-Mora, A., Bajic, V.B., Papatsenko, D.A., et al. (2018) HOCOMOCO: Towards a complete collection of transcription factor binding models for human and mouse via large-scale ChIP-Seq analysis. Nucleic Acids Res, 46, D252–D259.

47. Hänzelmann, S., Castelo, R. and Guinney, J. (2013) GSVA: Gene set variation analysis for microarray and RNA-Seq data. BMC Bioinformatics, 14, 1–15.

48. Federico, A. and Monti, S. (2020) hypeR: an R package for geneset enrichment workflows. Bioinformatics, 36, 1307–1308.

49. Giarratana, M.C., Rouard, H., Dumont, A., Kiger, L., Safeukui, I., Le Pennec, P.Y., François, S., Trugnan, G., Peyrard, T., Marie, T., et al. (2011) Proof of principle for transfusion of in vitro-generated red blood cells. Blood, 118, 5071–5079.

50. Sher, F., Hossain, M., Seruggia, D., Schoonenberg, V.A.C., Yao, Q., Cifani, P., Dassama, L.M.K., Cole, M.A., Ren, C., Vinjamur, D.S., et al. (2019) Rational targeting of a NuRD subcomplex guided by comprehensive in situ mutagenesis. Nat Genet, 51, 1149–1159.

51. Giarratana, M.C., Kobari, L., Lapillonne, H., Chalmers, D., Kiger, L., Cynober, T., Marden, M.C., Wajcman, H. and Douay, L. (2005) Ex vivo generation of fully mature human red blood cells from hematopoietic stem cells. Nature Biotechnology 2004 23:1, 23, 69–74.

52. Wangen, J.R., Eidenschink Brodersen, L., Stolk, T.T., Wells, D.A. and Loken, M.R. (2014) Assessment of normal erythropoiesis by flow cytometry: important considerations for specimen preparation. Int J Lab Hematol, 36, 184–196.

53. Siatecka, M. and Bieker, J.J. (2011) The multifunctional role of EKLF/KLF1 during erythropoiesis. Blood, 118, 2044.

54. Kalra, I.S., Alam, M.M., Choudhary, P.K. and Pace, B.S. (2011) Krüppel-like Factor 4 activates HBG gene expression in primary erythroid cells. Br J Haematol, 154, 248–259.

55. Xiong, Q., Zhang, Z., Qu, H., Ruan, X., Wang, H., Zhang, Q., Qi, H., Li, Y., Sun, H., Chang, K.-H., et al. (2012) Deciphering the Cis- and Trans-regulatory Roles of KLF6 in Primitive Hematopoiesis. Blood, 120, 4730.

56. Hänzelmann, S., Castelo, R. and Guinney, J. (2013) GSVA: Gene set variation analysis for microarray and RNA-Seq data. BMC Bioinformatics, 14, 1–15.

57. Kuleshov, M. V., Jones, M.R., Rouillard, A.D., Fernandez, N.F., Duan, Q., Wang, Z., Koplev, S., Jenkins, S.L., Jagodnik, K.M., Lachmann, A., et al. (2016) Enrichr: a comprehensive gene set enrichment analysis web server 2016 update. Nucleic Acids Res, 44, W90.

58. Evangelista, J.E., Clarke, D.J.B., Xie, Z., Lachmann, A., Jeon, M., Chen, K., Jagodnik, K.M., Jenkins, S.L., Kuleshov, M. V., Wojciechowicz, M.L., et al. (2022) SigCom LINCS: data and metadata search engine for a million gene expression signatures. Nucleic Acids Res, 50, W697–W709.

59. Wang, X.Y., Gelfanov, V., Sun, H. Bin, Tsai, S. and Yang, Y.C. (1999) Distinct actions of interleukin-9 and interleukin-4 on a hematopoietic stem cell line, EMLC1. Exp Hematol, 27, 139–146.

60. Kuhikar, R., Khan, N., Philip, J., Melinkeri, S., Kale, V. and Limaye, L. (2020) Transforming growth factor β1 accelerates and enhances in vitro red blood cell formation from hematopoietic stem cells by stimulating mitophagy. Stem Cell Res Ther, 11.

61. Zermati, Y., Fichelson, S., Valensi, F., Freyssinier, J.M., Rouyer-Fessard, P., Cramer, E., Guichard, J., Varet, B. and Hermine, O. (2000) Transforming growth factor inhibits erythropoiesis by blocking proliferation and accelerating differentiation of erythroid progenitors. Exp Hematol, 28, 885–894.

62. Rochira, V., Zirilli, L., Madeo, B., Maffei, L. and Carani, C. (2009) Testosterone action on erythropoiesis does not require its aromatization to estrogen: Insights from the testosterone and estrogen treatment of two aromatase-deficient men. J Steroid Biochem Mol Biol, 113, 189–194.

63. Azad, P., Villafuerte, F.C., Bermudez, D., Patel, G. and Haddad, G.G. (2021) Protective role of estrogen against excessive erythrocytosis in Monge’s disease. Experimental & Molecular Medicine 2021 53:1, 53, 125–135.

64. Bandach, I., Segev, Y. and Landau, D. (2021) Experimental modulation of Interleukin 1 shows its key role in chronic kidney disease progression and anemia. Scientific Reports 2021 11:1, 11, 1–14.

65. Ratajczak, A., Marlicz, W., Keidel, A., Machalinski, B., Ratajczak, M.Z. and Gewirtz, A.M. (1997) Effect of Interleukin-1 α and Interleukin-1 β on Erythroid Progenitor Cell Growth in Serum Free Cultures: An In Vitro Study Relevant to the Pathogenesis of the Anemia of Chronic Disease. Hematology, 2, 21–28.

66. Schooley, J.C., Kullgren, B. and Allison, A.C. (1987) Inhibition by interleukin-1 of the action of erythropoietin on erythroid precursors and its possible role in the pathogenesis of hypoplastic anaemias. Br J Haematol, 67, 11–17.

67. Mason, J.E., Nie, X., Coca, D. and Andrews, P.W. (2025) Stem cell fate decisions: Substates and attractors. Stem Cell Reports, 20, 102532.

68. Mead, B.E., Ordovas-Montanes, J., Braun, A.P., Levy, L.E., Bhargava, P., Szucs, M.J., Ammendolia, D.A., MacMullan, M.A., Yin, X., Hughes, T.K., et al. (2018) Harnessing single-cell genomics to improve the physiological fidelity of organoid-derived cell types. BMC Biology 2018 16:1, 16, 62-.

69. Huang, S., Eichler, G., Bar-Yam, Y. and Ingber, D.E. (2005) Cell Fates as High-Dimensional Attractor States of a Complex Gene Regulatory Network. Phys Rev Lett, 94, 128701.

70. Weng, C., Yu, F., Yang, D., Poeschla, M., Liggett, L.A., Jones, M.G., Qiu, X., Wahlster, L., Caulier, A., Hussmann, J.A., et al. (2024) Deciphering cell states and genealogies of human haematopoiesis. Nature 2024 627:8003, 627, 389–398.

71. To, K., Fei, L., Pett, J.P., Roberts, K., Blain, R., Polański, K., Li, T., Yayon, N., He, P., Xu, C., et al. (2024) A multi-omic atlas of human embryonic skeletal development. Nature 2024 635:8039, 635, 657–667.

72. Zuo, Z., Cheng, X., Ferdous, S., Shao, J., Li, J., Bao, Y., Li, J., Lu, J., Jacobo Lopez, A., Wohlschlegel, J., et al. (2024) Single cell dual-omic atlas of the human developing retina. Nature Communications 2024 15:1, 15, 1–21.

73. Ounadjela, J.R., Zhang, K., Kobayashi-Kirschvink, K.J., Jin, K., J. C. Russell, A., Lackner, A.I., Callahan, C., Viggiani, F., Dey, K.K., Jagadeesh, K., et al. (2024) Spatial multiomic landscape of the human placenta at molecular resolution. Nature Medicine 2024 30:12, 30, 3495–3508.

74. Ramírez, F., Ryan, D.P., Grüning, B., Bhardwaj, V., Kilpert, F., Richter, A.S., Heyne, S., Dündar, F. and Manke, T. (2016) deepTools2: a next generation web server for deep-sequencing data analysis. Nucleic Acids Res, 44, W160–W165.

75. Love, M.I., Huber, W. and Anders, S. (2014) Moderated estimation of fold change and dispersion for RNA-seq data with DESeq2. Genome Biol, 15, 1–21.

