## Supplementary Figures S1-S10 for "Directing stem cell differentiation by chromatin state approximation"

**Included in this document:**

Supplementary Figure S1 to S10

Supplementary References

**Not included in this document:**

Supplementary Table S1. Dataset overview with quality control metrics, principal component values, erythroid distance values, and chromatin module scores

Supplementary Table S2. Flow cytometry data

Supplementary Table S3. Mixed model coefficients

Supplementary Table S4. ATAC-seq peaks with AUC values

Supplementary Table S5. Gene set enrichment results in cell type-specific chromatin modules

Supplementary Table S6. Motif enrichments in cell type-specific chromatin modules

Supplementary Table S7. Motif enrichments in concordant/discordant regions

Supplementary Table S8. Perturbation gene set overlaps with transcription factors motifs that are enriched in discordant regions

Supplementary Table S9. Differentially accessible peaks in adapted protocols vs 5E

Supplementary Table S10. Materials used in this study

Supplementary Table S11. Oligonucleotides used in this study

### SUPPLEMENTARY FIGURES AND LEGENDS

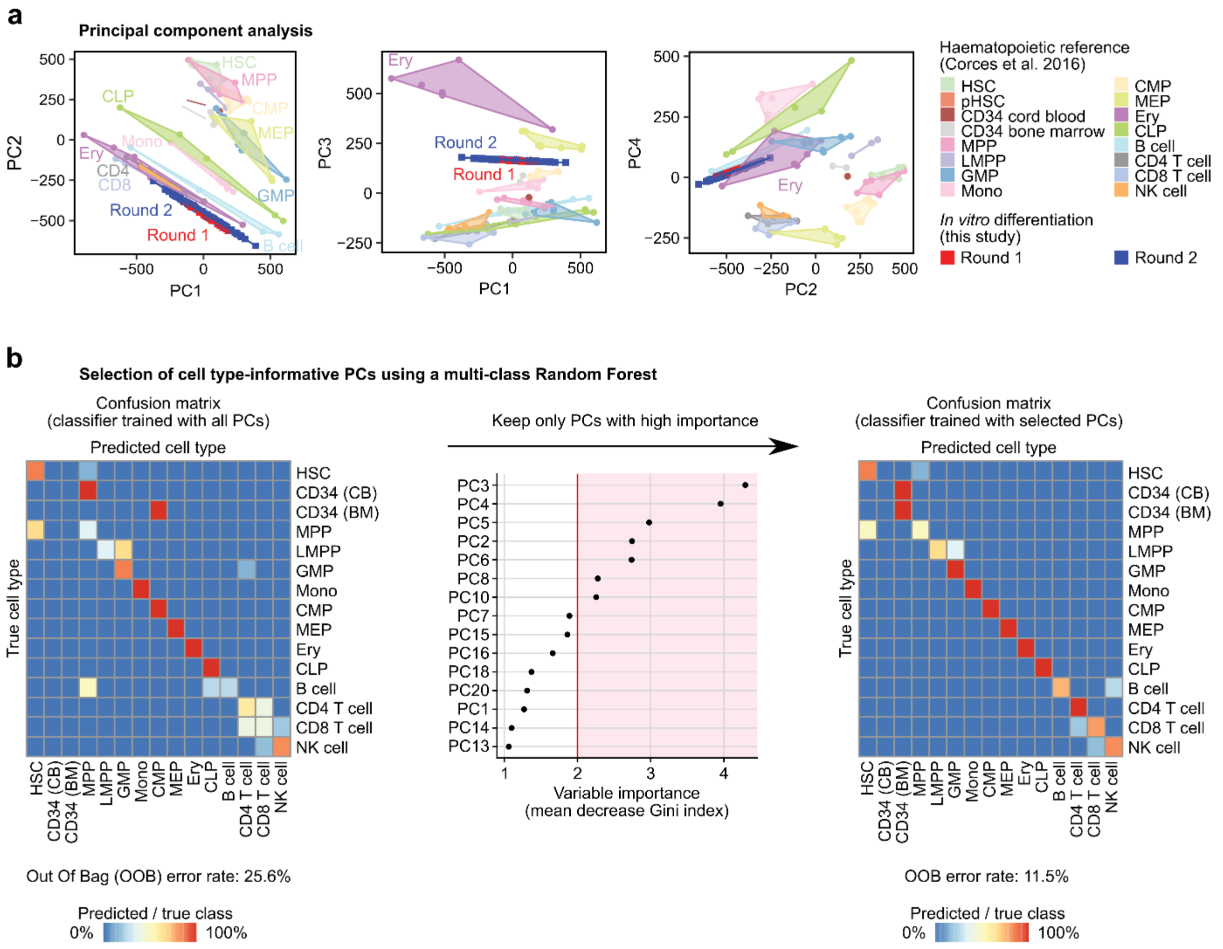


**Supplementary Figure S1. Extracting cell identity-relevant information from an ATAC-seq reference using Machine Learning (Random Forests).**

**a**, Principal component analysis of ATAC-seq samples of purified cell types (1) on all chromatin regions, visually depicting how some principal components (PCs) contain more cell identity information than others (e.g., PC1< PC3).

**b**, Confusion matrix of a multi-class Random Forest classifiers trained on the reference cell type data using PCs 1-78 (left). The most relevant PCs for classification (ranked by the mean decrease in Gini index, middle) were selected for a second Random Forest iteration (right), which reduced classification error by 58%. The selected, cell type-informative PCs were used for the PCA distance calculations in **Figure 1**.

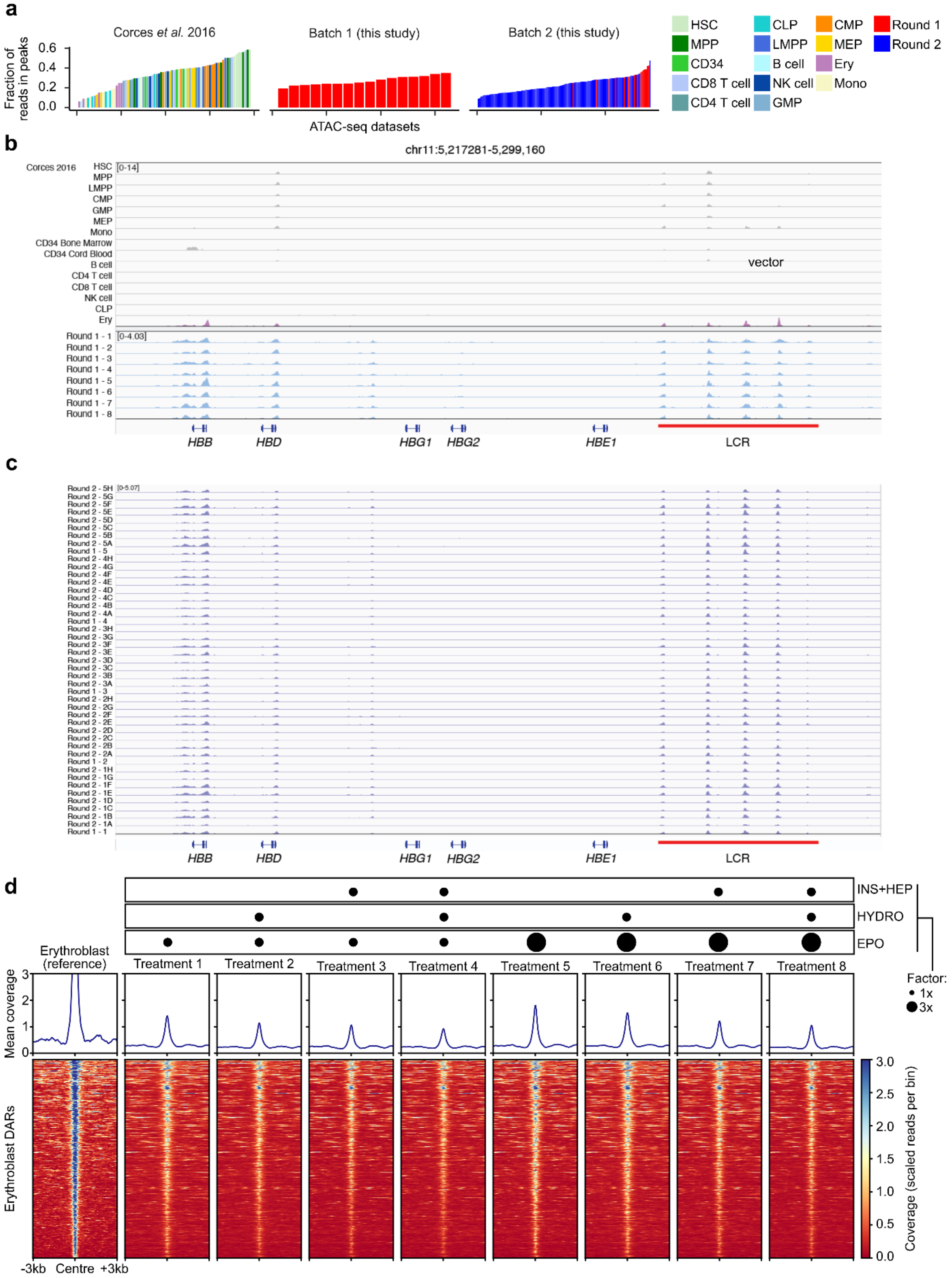


**Supplementary Figure S2. Quality control and example chromatin patterns of ATAC-seq from *in vitro*-differentiated erythroid cells.**

**a**, Fraction of Reads in Peaks (FRiP) scores for all ATAC-seq samples used in this study (**Supplementary Table S1**). Samples with a minimum score of 0.05 were kept. Scores were generated with MultiQC within the nf-core/atacseq pipeline.

**b,c** Visualisation of ATAC-seq tracks at the beta globin locus in reference datasets (Corces et al. 2016) and in cells from Round 1 (b) or Round 2 (c) of erythroid differentiation. Unlike embryonic (*HBE*) and foetal genes (HBG1/2), adult beta-like globin genes (*HBB*, *HBD*) and their 5' enhancers are accessible in erythroid cells. An interactive version of these data is available via the UCSC genome browser at https://genome.ucsc.edu/cgi-bin/hgTracks?hubUrl=http://fh.ccri.at/ML2Cell_trackhub/hub.txt, or by using a pre-configured IGV session at https://fh.ccri.at/ML2Cell_trackhub/igv_session_complete.xml.

**d**, Sorted region heatmaps comparing erythroblast-specific regions in FACS-sorted erythroblast samples(1) and from *in-vitro* samples of round 1 from this study. Each heatmap was generated using merged alignment files for all replicates from each condition (same regions as in **Fig. 1b**).

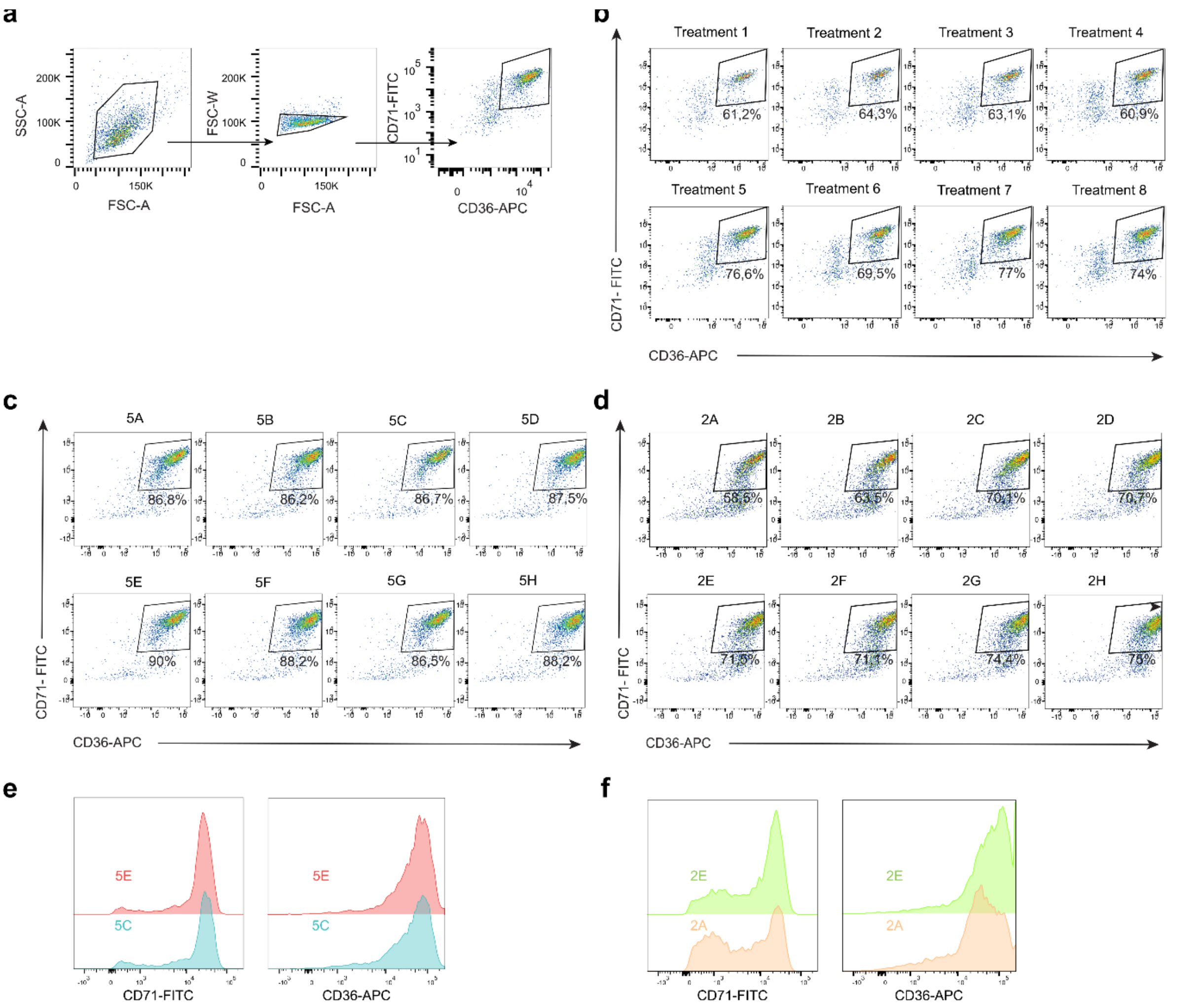


**Supplementary Figure S3. Flow cytometry gating strategy and example results after erythroid differentiation.**

**a**, Gating strategy used to select viable cell populations and erythroblastic, differentiated cells. Left: Forward scatter area (FSC-A) vs side scatter area (SSC-A); middle: Forward scatter width (FSC-W) vs forward scatter area (FSC-A); right: APC-A (anti-CD36) vs FITC-A (anti-CD71) to gate the double positive CD36+/CD71+ population.

**b,c,d,** CD36/CD71 expression by flow cytometry in cells of round-1 treatments (**b**), all round-2 treatments stemming from round-1 treatment 5 (5A-5H) (**c**), and all round-2 treatments stemming from round-1 treatment 2 (2A-2H) (**d**).

**e,f,** Marginal distributions for CD71 (FITC) and CD36 (APC) for two round-2 conditions stemming from round-1 treatment 5 (i.e., the best treatment of round 1) (**e**), including treatment 5E (the overall best) and another treatment. For comparison, samples stemming from round-1 treatment 2 are shown (**f**). The treatments shown (5C, 2E, 2A) were chosen for their characteristic differences in marker distribution.

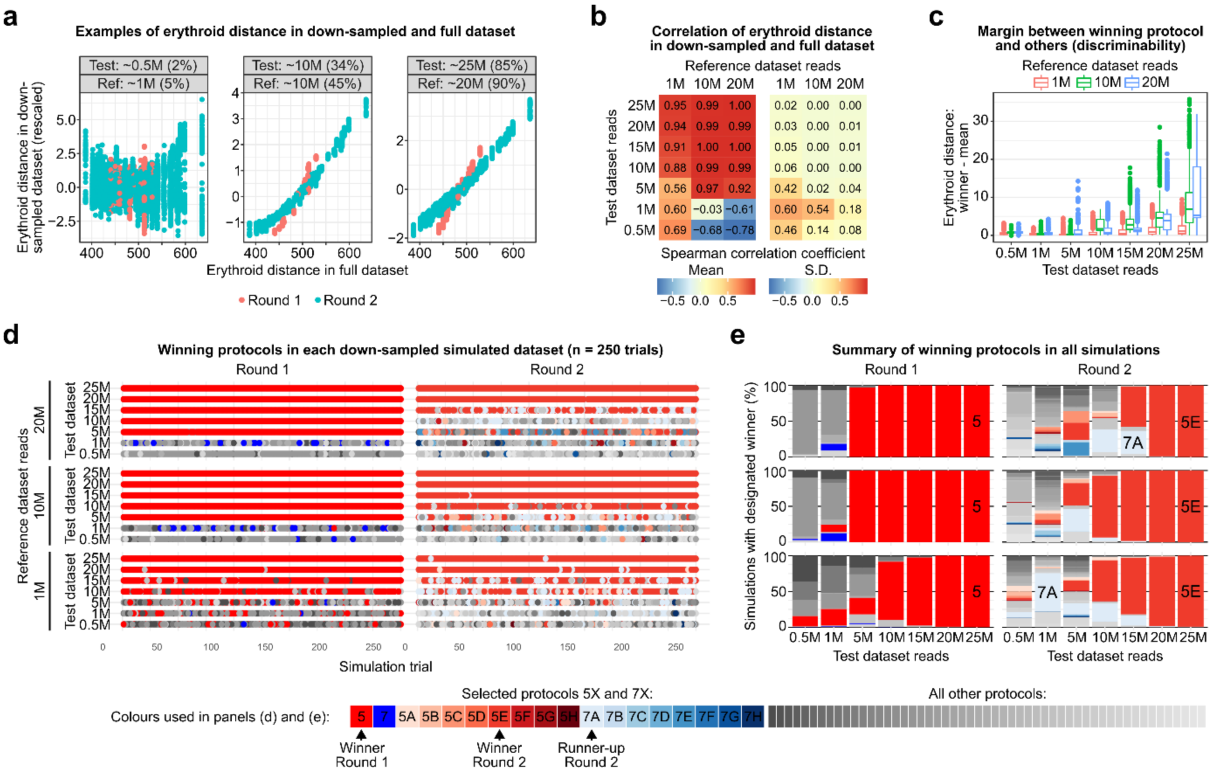


| | 1M | 10M | 20M | 1M | 10M | 20M |
|---|---|---|---|---|---|---|
| 25M | 0.95 | 0.99 | 1.00 | 0.02 | 0.00 | 0.00 |
| 20M | 0.94 | 0.99 | 0.99 | 0.03 | 0.00 | 0.01 |
| 15M | 0.91 | 1.00 | 1.00 | 0.05 | 0.00 | 0.01 |
| 10M | 0.88 | 0.99 | 0.99 | 0.06 | 0.00 | 0.00 |
| 5M | 0.56 | 0.97 | 0.92 | 0.42 | 0.02 | 0.04 |
| 1M | 0.60 | −0.03 | −0.61 | 0.60 | 0.54 | 0.18 |
| 0.5M | 0.69 | −0.68 | −0.78 | 0.46 | 0.14 | 0.08 |

**Supplementary Figure S4. Impact of sequencing depth on chromatin distance fidelity.**

Results of a down-sampling analysis on chromatin distance to erythroblasts. Read counts assigned to peaks in both the reference and test datasets were computationally downsampled (250 random trials) to defined percentages of the original read counts to simulate differences in sequencing depth.

**a,** Pairwise comparison of the erythroid distance in down-sampled dataset (y-axis) vs. the erythroid distance in the full dataset (x-axis, as reported in the paper) for three example read depths. The distance metric has been re-scaled per random trial to make the numbers comparable.

**b,** Heatmaps summarizing the mean (left) and standard deviation (S.D., right) of Spearman rank correlation coefficients for different sequencing depths in the test dataset (rows) vs reference dataset (columns). Down-sampled distances are highly correlated with the original distances at >10M reads, but at lower depths the correlation is lost. Even with just 1M reference reads, a good correlation is achieved if the test data is deep enough. The results are less stable (higher S.D.) with fewer reads.

**c,** Boxplots showing the margin (absolute difference) between the sample with the lowest erythroid distance and the average of all other erythroid distances in one random trial. The margin decreases notably with sequencing depth, meaning that the choice "best protocol" becomes increasingly random and based on small differences.

**d,** Plots showing the "winning" protocol in each of the 250 random trials at different read depths. The winners after round 1 and round 2 are shown separately (left and right, respectively). Blue colours indicate protocols 5X (including the best overall protocol 5E) and red colours indicates 7X (the runner-up). This confirms that the winning protocol has been chosen in most trials even at lower sequencing depths until <10M test reads, at which point the results become more diverse.

**e,** Stacked barplots showing the distribution of winning protocols across all trials (summary of the data in panel d).

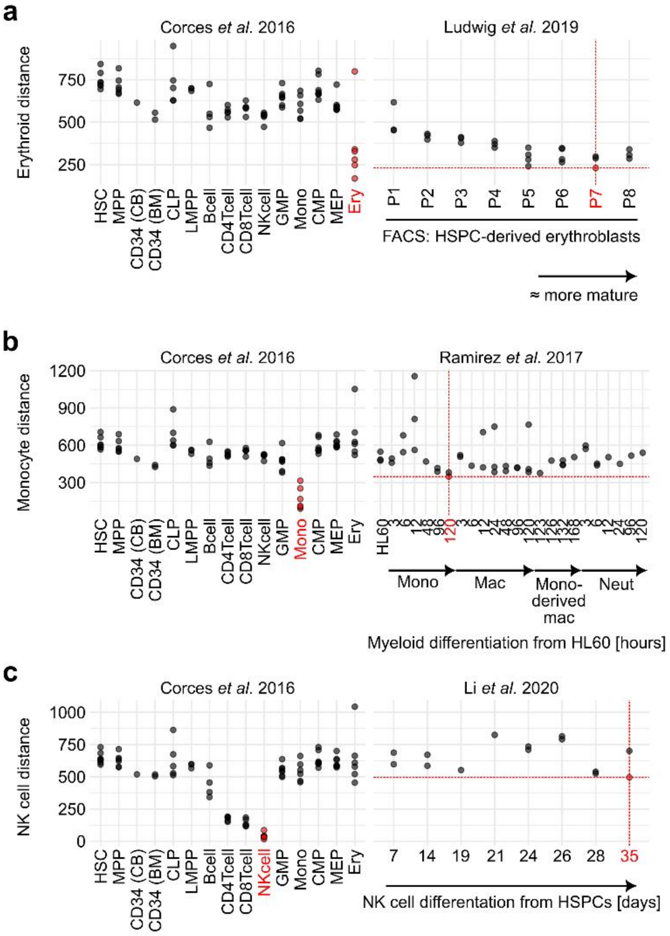


**Supplementary Figure S5. Chromatin distance discriminates top-performing in vitro samples in independent datasets and diverse targets.**

Comparison if independent datasets to defined haematopoietic target cell types. Analogously to the analyses shown in the main paper, chromatin distances were calculated with respect to the haematopoietic reference (2) using the chromatin distance in a joint PCA of all datasets. Samples in red corresponds to the condition with the sample of shortest chromatin distance in the panel. Left column: recalculation of the chromatin distances to the centroid of (a) Ery, (b) monocytes or (c) NK cells in the reference dataset as a benchmark. Right:

**a**, Chromatin distance of gated cell populations of *in-vitro*-differentiating erythroblasts with different markers generated by Ludwig *et al*. (3).

**b**, Chromatin distance to the monocyte centroid for samples of *in-vitro*-differentiating monocytes, macrophages, monocyte-derived macrophages, or neutrophils by Ramirez *et al*. (4).

**c**, Chromatin distance to the NK cell centroid for samples of in-vitro-differentiating NK cells by Li *et al.* (5).

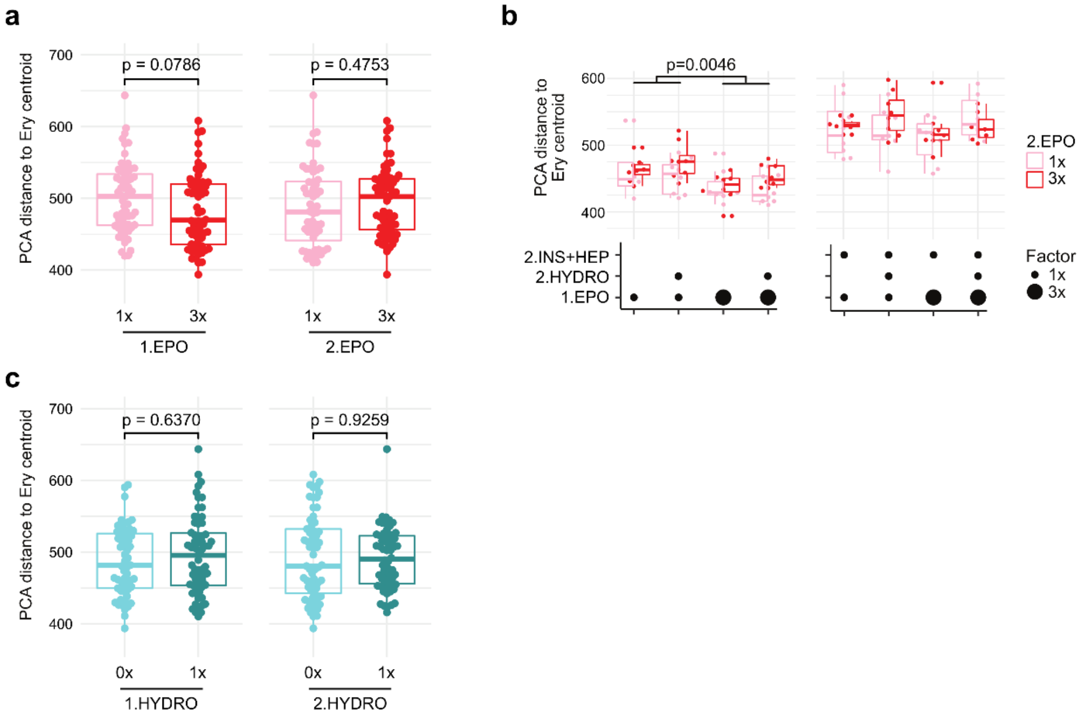


**Supplementary Figure S6. Time-sensitive and conditional contribution of ingredients on erythroid distance.**

**a,** Box plot showing the median, +/- interquartile range of the distance to the erythroblast centroid for samples of differentiation round 2 (n=119 samples), divided by whether they received EPO during round 1 (left) or 2 (right). All P values on the figure correspond to a two-sided Wilcoxon rank sum test and were adjusted for multiple hypothesis testing using the Benjamini-Hochberg method.

**b,** Same data as in a) but dividing the samples by whether they received INS+HEP at round 2 (right) or not (left), then by their exact combination of 1.EPO and 2.HYDRO, to emphasise the conditional EPO effects in the absence of 2.INS+HEP.

**c,** Effects of Hydrocortisone (HYDRO) at round 1 (left) and round 2 (right). Samples and statistical tests are the same as in panel **a.**

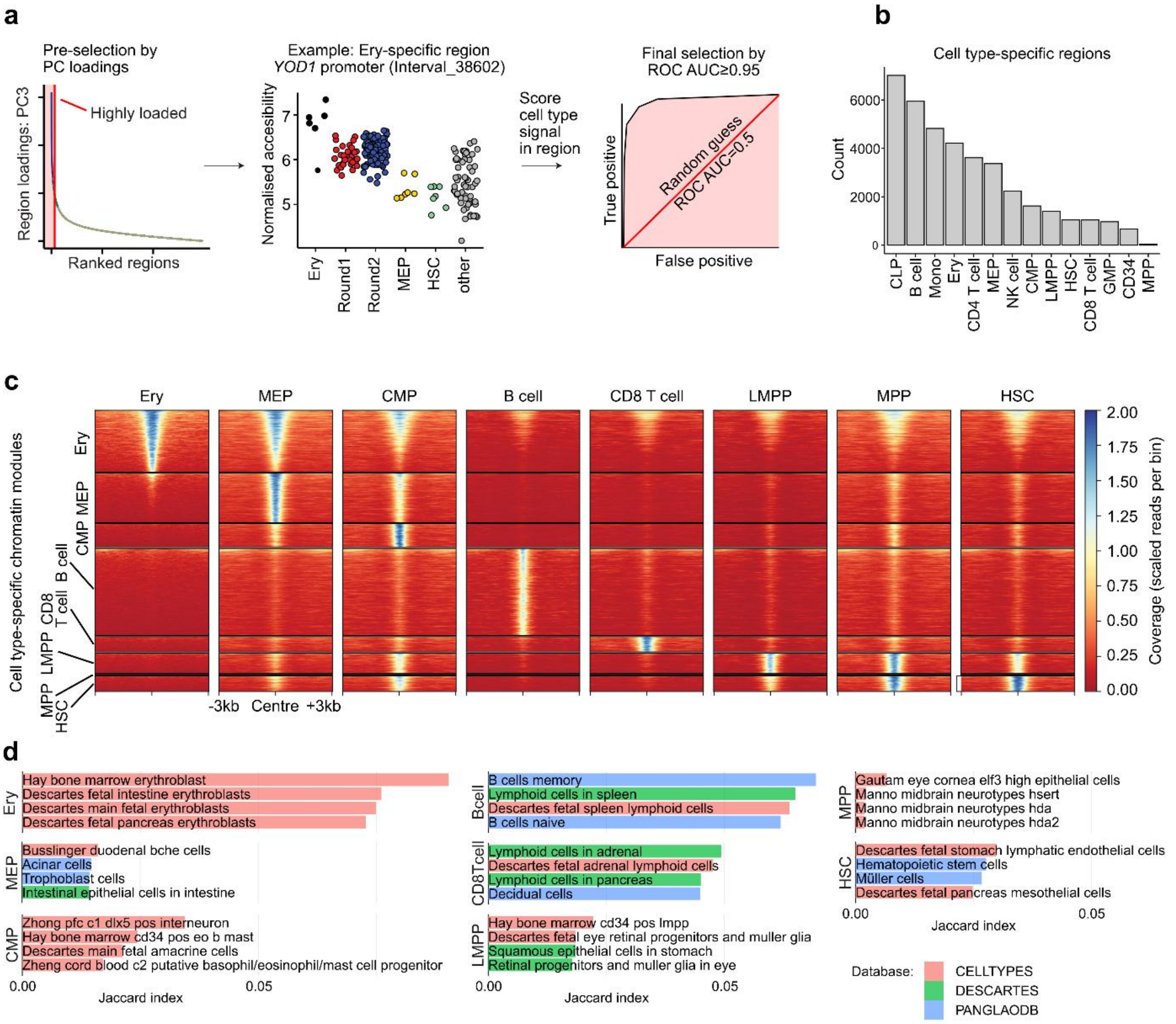


**Supplementary Figure S7. Strategy of discovery of cell identity peaks for each cell type.**

**a**, Workflow for selecting cell type-specific chromatin modules. Left: diagram depicting the loadings of all chromatin regions for an example principal component (specifically PC3). The most highly loaded peaks for each PC were shortlisted for receiver-operating characteristic (ROC) analysis. Centre: normalized ATAC-seq read counts at peak “interval_38602”, which overlaps with the *YOD1* promoter and is highly predictive of Ery cell type. Cell type-specific regions such as this one are the highest scoring for that cell type above all other samples, which makes the accessibility predictive of the cell type. Right: all regions with cell type-specific accessibility (like the *YOD1* promoter in the example) were selected by assessing the ROC area under the curve (AUC). A steep incline in the ROC curve and high AUC value indicates strong discriminative power.

**b**, Amount of cell type-specific regions detected for each cell type after assessing 355,314 regions.

**c**, Chromatin accessibility of the cell identity chromatin modules (rows) in the haematopoietic reference dataset (1) (all samples of the same cell type merged).

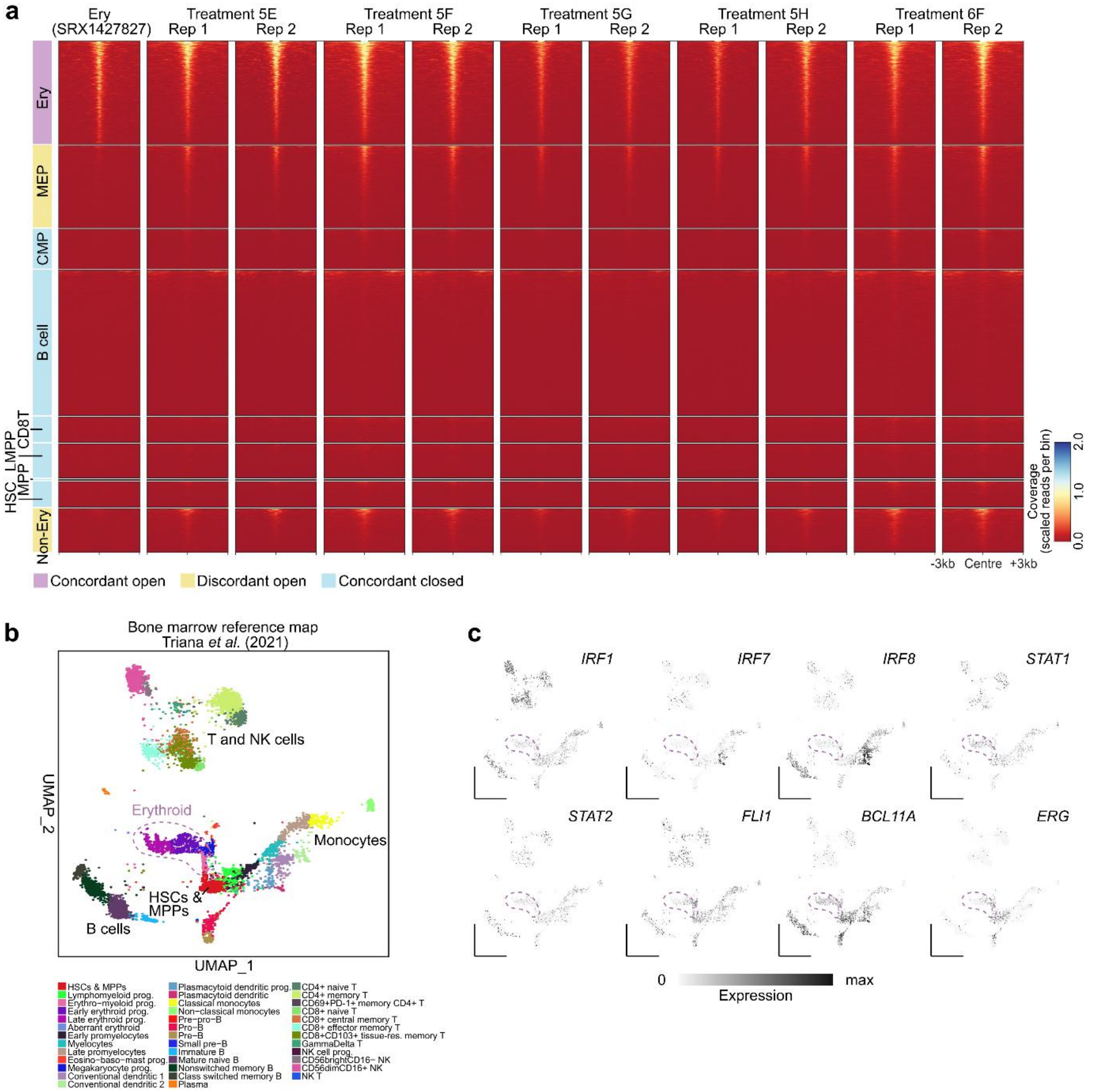


**Supplementary Figure S8. Concordant and discordant region sets and expression of priority TFs.**

**a,** Coverage of ATAC-seq reads in cell type-specific chromatin modules for a representative Ery reference samples and *in-vitro* samples, including samples from treatment 5E for comparison (the overall best treatment). Colours of each module indicate whether the module is concordant open, concordant closed, or discordant. These groups have been selected based on the values in the overall best sample (5E) and the reference erythroblasts.

**b**, UMAP showing whole transcriptomes (plus 97 surface markers) of haematopoietic cells derived from the bone marrow by Triana *et al.* (6). Pre-processed data from the original publication was used without further modification.

**c**, Expression levels of selected priority TFs (whose binding sites are overly abundant in discordant regions) from **Figure 4**. The window containing erythroid cells has been highlight manually with dashed lines to aid interpretation. Many priority TFs are expressed outside the erythroid lineage.

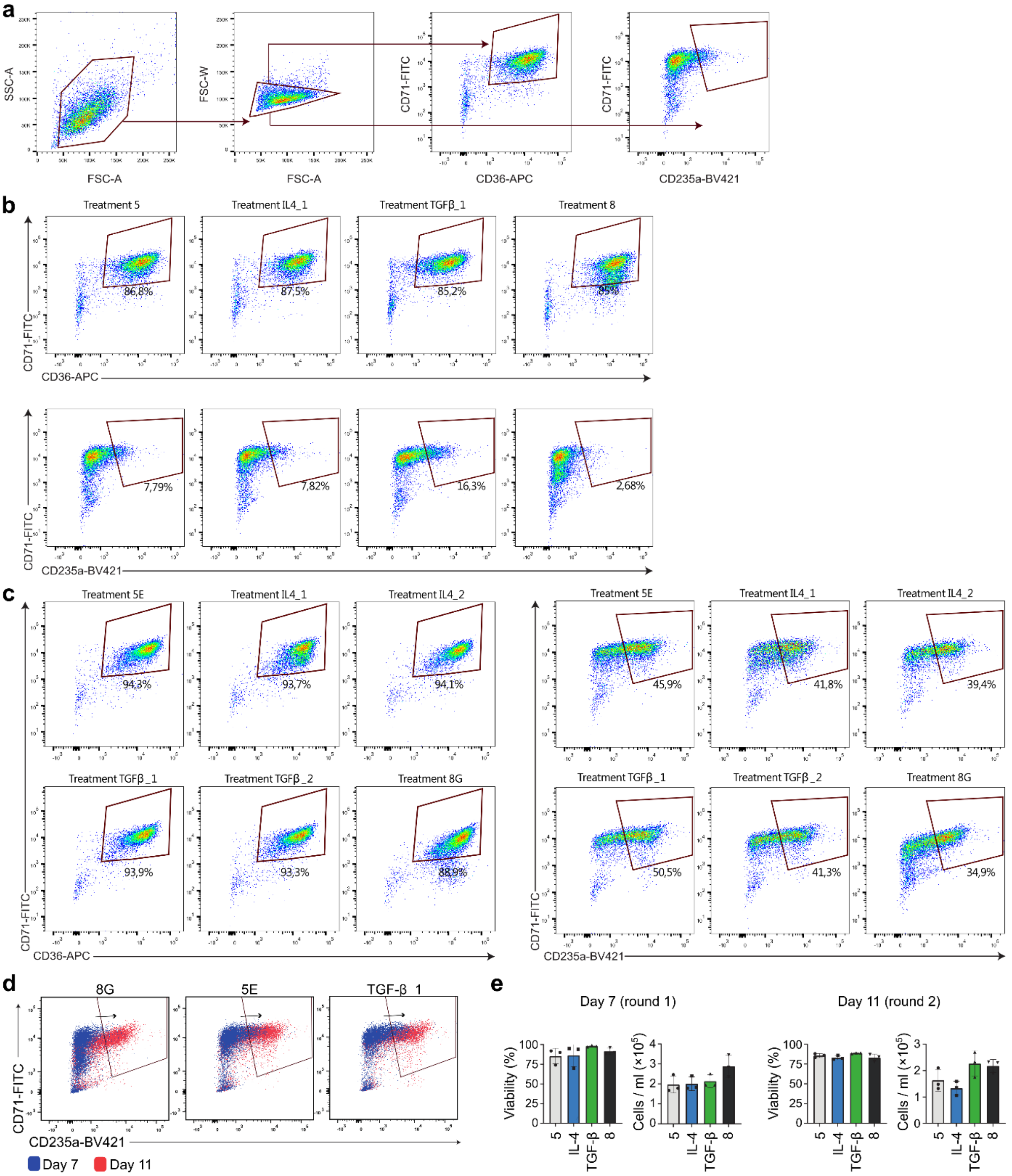


### Supplementary Figure S9. Flow cytometry and viability analysis.

**a**, Gating strategy used to select viable cell populations and erythroblastic, differentiated cells. Left: Forward scatter area (FSC-A) vs side scatter area (SSC-A); middle: Forward scatter width (FSC-W) vs forward scatter area ( FSC-A); right: APC-A (anti-CD36) vs FITC-A (anti-CD71) to gate the double positive CD36+/CD71+ population and FITC-A (anti-CD71) vs BV241 (anti CD235a) to gate the double positive CD71+/CD235a+ population.

**b**, CD36+/CD71+ expression (top) and CD71+/CD235a+ expression (bottom) by flow cytometry in cells of round-1 (day 7) of differentiation.

**c**, CD36+/CD71+ expression (top) and CD71+/CD235a+ expression (bottom) by flow cytometry in cells of round 2 (day 11) of differentiation.

**d,** Mean fluorescence intensity (MFI) plots indicating the distribution of CD71 and CD235a expression at day 7 (blue) and day 11 (red) using flow cytometry.

**e,** Viability and number of cells at day 7 (left) and day 11 (right) of erythroid differentiation, comparing protocols 8G, 5E, and 5E supplemented with IL-4 or TFG-β.

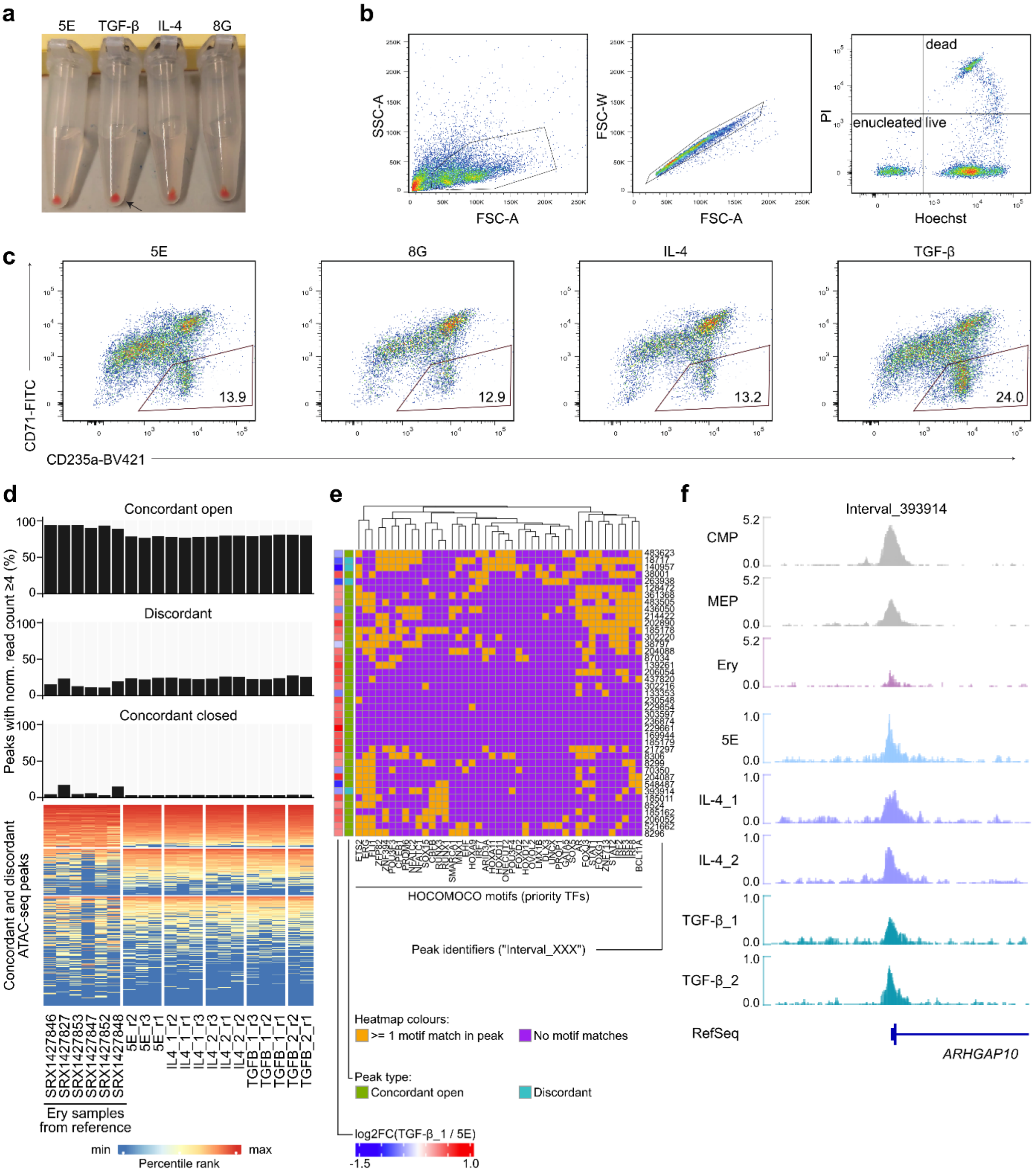

**Supplementary Figure S10. Functional and chromatin evaluation of modified treatments.**

**a**, Representative cell pellets at day 14 of differentiation.

**b**, Gating strategy used to select enucleated cells. Left: Forward scatter area (FSC-A) vs side scatter area (SSC-A); middle: Forward scatter width (FSC-W) vs forward scatter area (FSC-A); right: PE-Texas Red (Propidium Iodide) vs Brilliant Ultraviolet 496 (Hoechst) to gate the double negative PI-/ Hoechst- particles indicating enucleated cells.

**c**, Representative gating CD235a-positive, CD71-negative cells at day 14, plotted in Fig. 5e.

**d**, Top: Barplots indicating the percentage of concordant open, discordant, and concordant closed peaks that were accessible (VST-normalized read count > 4) in erythroblast reference samples (sample names starting with "SRX") compared to *in-vitro*-differentiated cells. Bottom: Percentile rank of each peak within a sample according to treatment and group (for the same samples as in the barplots). Peaks belonging to the concordance groups were assigned a percentile accessibility rank, then seriated per group to emphasise altered ranking across treatments. Percentiles below 40% are set to blue.

**e**, Heatmap indicating the presence of binding sites of priority TFs (matches to the DNA sequence motif from the HOCOMOCO database) for differentially accessible peaks between treatments TGF-β_1 and 5E. Orange squares indicate at least one motif match was present. The colour bars on the left indicate the $log_2$ fold change and the peak group. The numbers in each row of the heatmap denoted the identified of the peaks as also shown in **Supplementary Table S9**.

f, Visualisation of ATAC-seq coverage at the promoter of *ARHGAP10* showing an example (Interval_393914) of a differentially accessible discordant peak whose accessibility is low in Ery, and reduced in TGF-β_1 compared to treatment 5E.

### SUPPLEMENTARY REFERENCES


1. Corces,M.R., Buenrostro,J.D., Wu,B., Greenside,P.G., Chan,S.M., Koenig,J.L., Snyder,M.P., Pritchard,J.K., Kundaje,A., Greenleaf,W.J., *et al.* (2016) Lineage-specific and single-cell chromatin accessibility charts human hematopoiesis and leukemia evolution. *Nature Genetics 2016 48:10*, 48, 1193–1203.

2. Corces,M.R., Buenrostro,J.D., Wu,B., Greenside,P.G., Chan,S.M., Koenig,J.L., Snyder,M.P., Pritchard,J.K., Kundaje,A., Greenleaf,W.J., *et al.* (2016) Lineage-specific and single-cell chromatin accessibility charts human hematopoiesis and leukemia evolution. *Nat Genet*, 48, 1193–1203.

3. Ludwig,L.S., Lareau,C.A., Bao,E.L., Nandakumar,S.K., Muus,C., Ulirsch,J.C., Chowdhary,K., Buenrostro,J.D., Mohandas,N., An,X., *et al.* (2019) Transcriptional States and Chromatin Accessibility Underlying Human Erythropoiesis. *Cell Rep*, 27, 3228-3240.e7.

4. Ramirez,R.N., El-Ali,N.C., Mager,M.A., Wyman,D., Conesa,A. and Mortazavi,A. (2017) Dynamic Gene Regulatory Networks of Human Myeloid Differentiation. *Cell Syst*, 4, 416-429.e3.

5. Li,K., Wu,Y., Li,Y., Yu,Q., Tian,Z., Wei,H. and Qu,K. (2020) Landscape and Dynamics of the Transcriptional Regulatory Network During Natural Killer Cell Differentiation. *Genomics Proteomics Bioinformatics*, 18, 501–515.

6. Triana,S., Vonficht,D., Jopp-Saile,L., Raffel,S., Lutz,R., Leonce,D., Antes,M., Hernández-Malmierca,P., Ordoñez-Rueda,D., Ramasz,B., *et al.* (2021) Single-cell proteo-genomic reference maps of the hematopoietic system enable the purification and massive profiling of precisely defined cell states. *Nature Immunology 2021 22:12*, 22, 1577–1589.
